# Current and future relevance of marine spatial planning for the distribution of epibenthic invertebrate and fish species in a heavily used regional sea

**DOI:** 10.1101/2025.02.10.637456

**Authors:** W. Nikolaus Probst, Jennifer Rehren, Casper Kraan, Holger Haslob, Hermann Neumann, Carsten Lemmen, Shubham Krishna, Kai Wirtz, Vanessa Stelzenmüller

**Affiliations:** Thünen-Institute of Sea Fisheries; Herwigstraße 31, 27572 Bremerhaven, Germany; Helmholtz-Zentrum Hereon, Institute of Coastal Systems – Analysis and Modeling, Max-Planck-Straße 1, 21502 Geesthacht

**Keywords:** species distribution models, North Sea, anthropogenic pressures, global warming, offshore wind farms, marine protected areas

## Abstract

The SNS has recently become a European hub for installations of offshore wind farms (OWF), while extensive areas have been designated as marine protected areas (MPAs). Together with the already noticeable effects of climate warming, the region transforms from an area dominated by free ranging fisheries and shipping into an industrial landscape dominated by stationary activities. To inform decision making processes around the spatial allocation of fisheries, conservation measures and licence areas for offshore renewables in the southern North Sea, we modelled the spatial distribution of 179 epibenthic invertebrate and fish species. We identify both current and future hotspots of epibenthic and demersal fish diversity and their overlap with OWFs and MPAs. Hotspots of epibenthic and fish diversity are mostly found along the English Coast, located within an extensive network of MPA and OWF sites, which may provide opportunities for conservation and fisheries alike. In the central SNS, the MPA network could be complemented by OWF, where co-use regulations with other human uses are excluded, to protect sensitive species as well as epibenthic and demersal fish communities. Scenarios for three different time periods (until 2040, 2070 & 2100) revealed that global warming might cause an increase in the probability of occurrence for more than half of the analysed demersal fish species (n = 63). The applications in our study demonstrate the relevance of comprehensive knowledge about current, near and far future distribution patterns of species and species communities to enhance the effectiveness of marine spatial planning including spatial conservation measures.

## Introduction

The southern North Sea (SNS) has been subject to human use for centuries, and plays a central role in the economic and cultural development of the surrounding regions. Its waters have supported thriving fisheries, highly frequented shipping routes, the exploration of sand, oil, and gas and numerous other anthropogenic activities (Emeis *et al*. 2015). The productivity of the SNS has been a cornerstone for sustaining coastal communities and a significant driver of economic growth (Halpern *et al*. 2012; OSPAR 2023). However, the ecosystems of the SNS are subject to multiple stressors ranging from climate change and intensive fishing to the expansion of maritime industries (McLean *et al*. 2019; Stelzenmüller *et al*. 2024). Offshore wind farms (OWF), are planned to increase substantially to ∼ 300 GW by 2050 to cover the increasing demand for renewable energy, and by then might occupy large areas in the SNS (Gusatu *et al*. 2020; Stelzenmüller *et al*. 2022; Li *et al*. 2023). In parallel, the exclusion of fisheries within marine protected areas (MPA) will expanded under the EU Biodiversity Strategy 2030 (EEA 2015; EC 2021; EC 2023; OSPAR 2023).

The combination of OWF and MPA will lead to closures for fishing with trawled gears, altering future impacts of fishing on habitats, the epibenthic fauna, and the demersal fish community of the SNS (Beauchard *et al*. 2017; Stelzenmüller *et al*. 2022; Bonsu *et al*. 2024). Bottom trawling is one of the main human activities in the SNS, targeting brown shrimp *Crangon crangon,* flatfish species such as plaice *Pleuronectes platessa* and sole *Solea solea* and roundfish such as haddoc*k Melanogrammus aeglefinus,*and cod *Gadus morhua.* Satellite based data on annual fishing effort suggest that some areas are trawled more than 10 times per year (ICES 2021). For conservation purposes it is therefore important to understand the distribution of threatened species, species of commercial interest and hotspots of species richness (Neumann *et al*. 2017; Probst *et al*. 2021; Weinert *et al*. 2021).

We here combined the data of three fisheries surveys covering the years 2014 to 2023 to compute 179 species distribution models (SDM) for epibenthic invertebrates and fish in the SNS to determine the degree of spatial overlap between core areas of species diversity and areas excluding trawled fisheries (OWFs and MPAs). We further explored the response of individual species to a suite of pressures such as fishing effort, temperature and distance to offshore wind farms.

## Materials & methods

### Data compilation and processing

Averaged monthly means over the time period 2004—2012 of dissolved oxygen, maximum shear stress, particulate organic carbon, temperature, zooplankton carbon at the sea floor, dissolved phosphate and nitrate in the water column, light attenuation and sea surface salinity (Figure S2, Table S2) were obtained from a high resolution model simulation(Wirtz *et al*. 2024), where the ecosystem model MAECS (Wirtz 2019) was coupled to the hydrodynamic model GETM and other earth system modular components such as benthic biogeochemistry through the coupling framework MOSSCO (Lemmen *et al*. 2018). For three future time periods (2030-2040, 2060-2070 and 2090-2100) data on temperature anomalies relative to 2004-2012 were obtained from a simulation with the coupled REMO-MPIOM-HAMOCC model (Mathis, Elizalde & Mikolajewicz 2019, their RCP 8.5 scenario ensemble member 174) to recalculate projected maps of sea bottom temperature and temperature range at the sea bottom (supplements S3).

Spatial information on installed and projected OWF (up to and including 2042) was obtained from 4Coffshore Ltd. (www.4coffshore.com). Data was filtered to exclude OWF marked with the status “cancelled”, “failed proposal” or “dormant”. For the calculation of the distance to offshore wind farms, used as a predictor for the SDMs, the data was filtered for OWF that were installed until 2022. For the overlay analysis all installed and projected (until 2042) OWF were combined into a unified shapefile. Fisheries intensity, expressed as swept area ratios (SAR), was downloaded from the International Council for the Exploration of the Sea (ICES) including subsurface and surface SAR averaged from 2009 until 2020 (ICES 2021). MPA shapefiles were downloaded from the World Database on Protected Areas provided by the UN Environmental Programme and the International Union for Conservation of Nature (UNEP-WCMC & IUCN 2024, www.protectedplanet.net).

All predictor variables were transformed to raster format on a rectangular raster grid with a resolution of 0.025° longitude x 0.025° latitude, which corresponds to a rectangle roughly 1.6 km x 2.8 km, or an average distance between grid cell centres of about 3 km. We also tested for collinearity requiring correlations R < 0.7 (Dormann *et al*. 2013). The size and extent of the spatial grid in this analysis was constrained by the spatial extent of the GETM SNS model domain ranging from −1°W to 9°E longitude and 51°N to 56°N latitude.

Fish and epibenthic occurrence and biomass data were obtained from (i) the Demersal Young Fish Survey DYFS, (ii) the North Sea International Bottom Trawl Survey NS-IBTS, and (iii) the beam trawl survey BTS (Figure 1). The three surveys were conducted in the SNS between 2014 and 2023 using beam or otter board trawls (Table 1).

**Figure 1.**
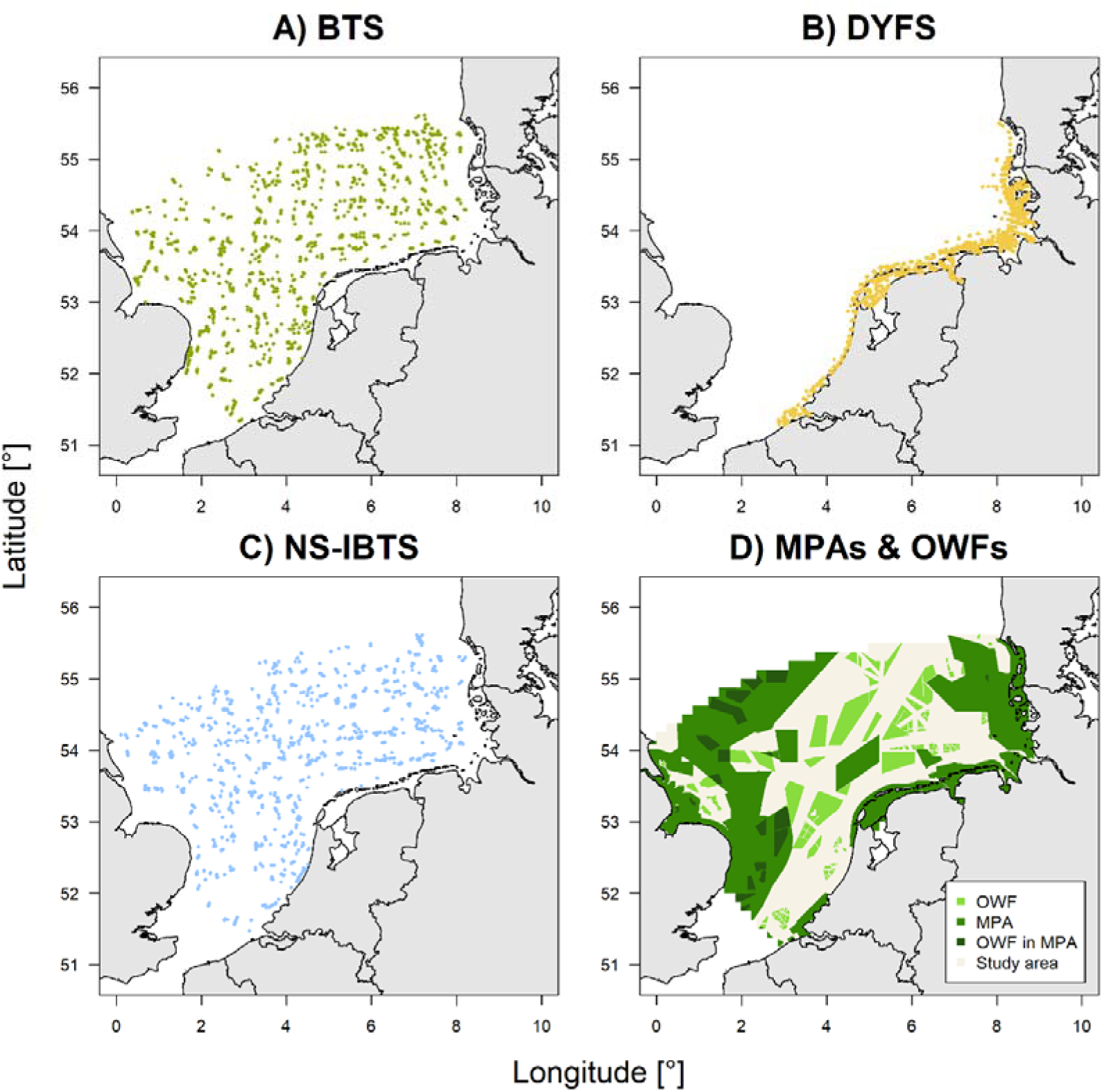
Location of samples (fishing hauls) from three fisheries surveys in the southern North Sea conducted between 2014 and 2023. A) Beam trawl survey BTS, B) Demersal Young Fish Survey DYFS, C) North Sea International Bottom Trawl Survey NS-IBTS; all available via the ICES DATRAS data portal). For details see Table 1. D) Location of offshore windfarms (OWF) and marine protected areas (MPA) in the study area.

**Table 1.**
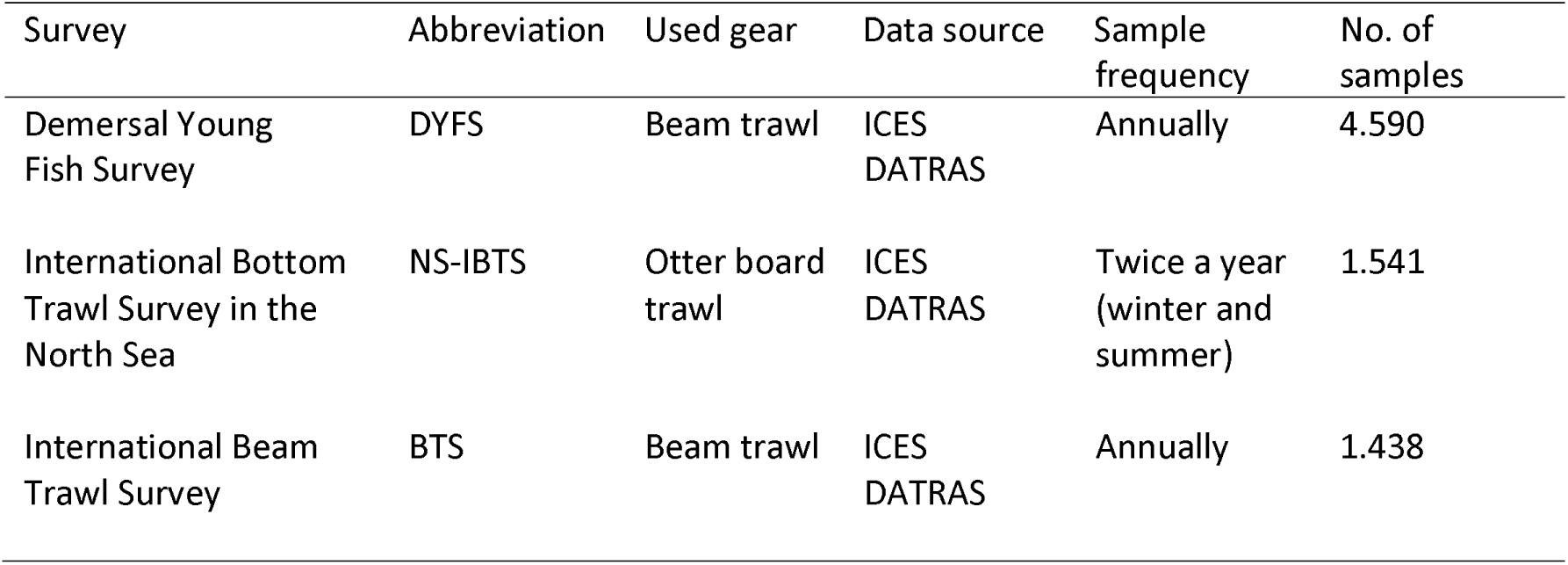
Overview of survey data from 2014 to 2023 used to model distributions of 180 epibenthic and fish species.

| Survey | Abbreviation | Used gear | Data source | Sample frequency | No. of samples |
| --- | --- | --- | --- | --- | --- |
| Demersal Young Fish Survey | DYFS | Beam trawl | ICES DATRAS | Annually | 4.590 |
| International Bottom Trawl Survey in the North Sea | NS-IBTS | Otter board trawl | ICES DATRAS | Twice a year (winter and summer) | 1.541 |
| International Beam Trawl Survey | BTS | Beam trawl | ICES DATRAS | Annually | 1.438 |

All surveys are coordinated by ICES and the data including records on haul length and survey information are available through the ICES DATRAS data portal (https://www.ices.dk/data/data-portals/Pages/DATRAS.aspx). As DATRAS only provides information on catch-per-hour, data on wing spread and towed distance were used to estimate densities (observed number of individuals per km²) and biomass (kg per km²). To fill partial data gaps on wing spread for otter board trawls, a regression between wingspread and depth was used (wing spread = 8.190 + 2.906 * log (Depth [m]), R²=0.170). For beam trawl gears, wing spread was set equal to beam width. If records lacked towed distances, the latter was estimated via regression from haul duration and towing speed. Partially missing values for towed distance were estimated by estimating the distance between the shoot and haul position using the function ‘distGeo’ from the r-package ‘geosphere’ (version 1.5-18). Samples without occurrence of any given species were considered as zero occurrence.

### Selection of species

We identified 179 fish and epibenthic invertebrate species that were recorded in at least 20 samples (i.e. fishing hauls) between 2014 and 2023 (Table S1). Even though the environmental data covered a time period from 2004 to 2012 (see supplements Table S2), we chose to use survey data from the most recent decade (2014 – 2023) to get an actual estimate of the species’ distributions, assuming patterns in oceanographical predictors from 2004 2012 remained comparable to the period of 2014 2023.

For fish species and several invertebrates including three commercially relevant crustacean taxa (brown shrimp *Crangon crangon*, brown crab *Cancer pagurus*, and Norway lobster *Nephrops norvegicus*) the biomass density in kg per km² was estimated using length-weight parameters (Weight[g] = a*Length [cm] ^ b) from FishBase (www.fishbase.org, data download at 08.02.2024 via the r-package ‘rfishbase’ using the ‘length-weight’-function), which were further complemented by an internal Thünen-data base (Wilhelms 2013). Length-weight parameters for European lobster (*Homarus Gammarus*) were obtained from (Pavičić *et al*. 2021). All parameters are included in supplementary material S1.

To address the objectives of this study, we defined different suites of species containing i) all modelled species, ii) epibenthic invertebrates, iii) demersal fish, iv) species of conservation concern and v) species of commercial relevance. Species of conservation concern were identified as species that were listed by the regional European IUCN list as “near threatened”, “vulnerable”, “endangered” or “critically endangered”, listed under the OSPAR red list of species, listed in Annex II of the EU Habitats Directive or were considered of national relevance in Germany (i.e. Norway lobster *Nephrops norvegicus*, European lobster *Homarus gammerus*, and the mud lobster *Upogebia deltaura*). Relevant commercial species were defined based on the expert knowledge of the authors.

### Modelling and evaluating species distributions and responses

For each species, the probability of occurrence (P_occ_) and the biomass density (kg per km²) were estimated using random forests (RF, Breiman 2001) implemented in the R-package ‘randomForest’ (version 4.7-1.1). Among other tested SDM methods, RF were found to be the most versatile algorithms allowing to include dependent and predictor variables of all types (i.e. continuous, categorical, nominal variables). The RF for species distribution (SDM) were used with default parameters except for setting the parameter ‘nodesize’ = 10, ‘maxnodes’ = 5 and ntree = 1000 to prevent overfitting (Probst, Wright & Boulesteix 2019). To account for the effects of different fishing gears used in the surveys, the gear category (small beam trawl with a width of 3 – 4 m, large beam trawl with a width of 6 – 8 m and otter board trawls) was included as a predictor variable. To account for spatial autocorrelation, the local Moran’s I for occurrence and biomass per km² were calculated (Anselin 1995) and included as a predictor in the RF (Georgian, Anderson & Rowden 2019).

To improve model performance, biomass records exceeding the 99.9% quantile of all biomass data were excluded from the training data set. Such outliers, however, were not removed for taxa with less than 50 occurrences. Thereby also very rare taxa such as sea horses *Hippocampus spp* were modelled and even though some authors suggest to consider only species with at least 50 occurrences (Virgili *et al*. 2018) we decided to implement a lowered cut-off threshold (n ≥ 20) to include rare species, which often are of conservation concern.

SDMs were used to predict spatial distributions in four time slices (2014–2023, 2030–2040, 2060–2070 and 2090–2100), which differed only with respect to sea bottom temperature and sea bottom temperature range (supplements S3). For each simulation, the data set was randomly divided into a training (75%) and validation data set (25%).

As evaluation metrics for SDMs which modelled P_occ_, we used the area-under-the-curve (AUC), true skill statistic (TSS) and the mean average error (MAE). For SDMs modelling biomass, Pearson’s product moment correlation between estimated values of biomass by the model vs. values derived from the validation data set was calculated. However, we did not define any threshold for AUC, TSS or MAE (for P_occ_) or the Pearson’s R (for biomass) to separate adequate from non-adequate models. Instead, we used all models from all species to identify core areas and hotspots considering each model as the best estimate of the true distribution. For completeness, however, all parameters are shown in supplements S1.

RF models allow to extract partial dependencies which show the relationship between single predictors and the response variable. The partial dependencies are given as x-y data and were regressed via a generalized linear model (GLM) to determine whether the slope of the partial dependency was significantly positive or negative. In these cases, the response was attributed as ‘increasing’ or ‘decreasing’, otherwise the response was classified as ‘neutral’.

### Defining core areas and demersal communities

SDM-based P_occ_ was used to identify core areas (CA) of species’ distribution. CA were calculated as absolute and relative core areas for each species. Absolute CA were areas in which P_occ_ > 0.8 and hence not each species had an absolute CA, as the majority of maximum P_occ_ was < 0.8. Relative CA were based on the 80%-quantile of all P_occ_-values indicating areas where species were found with the highest probability. Relative P_occ_-CA could be identified for each species and accordingly relative CA were also calculated based on biomass SDMs. Contrary to P_occ_, absolute CA based on biomass could not be calculated because of the scale difference of absolute biomass values between species (P_occ_ always ranges between 0 and 1 whereas biomass per km² can range between 0 and - the maximum observed value of the species).

A matrix of P_occ_ values from 63 demersal fish species was used to identify community clusters using K-means clustering (Hartigan & Wong 1979). To identify the optimal number of clusters a scree plot was implemented to visually evaluate which numbers of clusters would provide the best information gain with as few clusters as possible by looking for a ‘knee’ in the scree plot determined by a 66 % reduction of the unclustered (K = 1) sum of squares error (supplements S5).

### Spatial overlap analysis

The relative P_occ_-CA of all species were overlaid with locations from OWF and MPA to identify regions of overlap. The percentage of overlap between MPAs and CA, OWFs and CA as well as the proportion of CA that did not overlap with any MPA or OWF were calculated by summing the area of all grid cells that overlapped with the according shapefiles (see Figure 4 for an example and Table 2 for summary of overlap statistics).

**Table 2.**
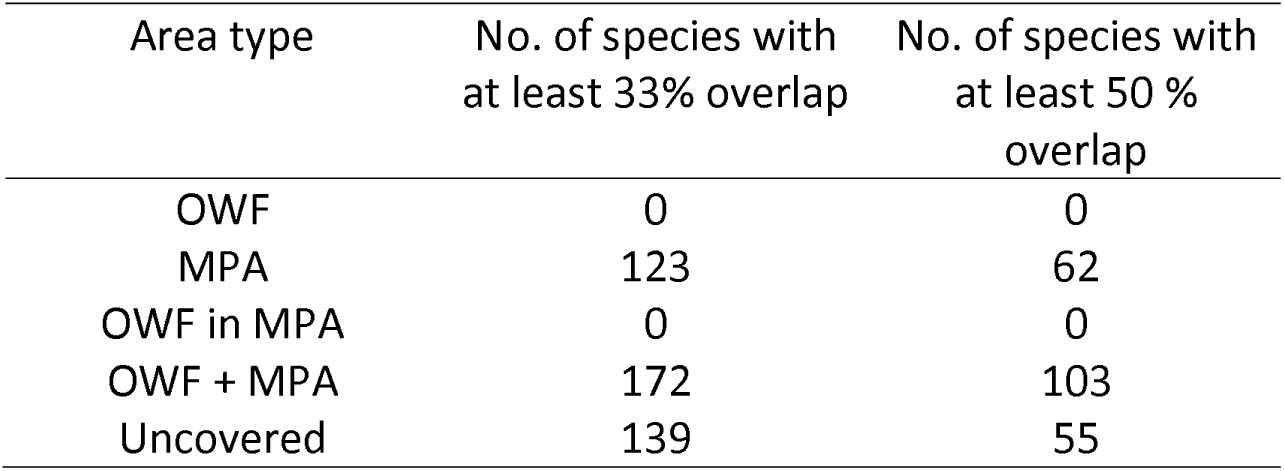
Number of species with 33% or 50 % overlap with offshore windfarms (OWF), marine protected areas (MPA), areas designated as both (OWF & MPA) and that are neither designated as OWFs nor MPAs. Total no. of species n = 179.

designated as OWFs nor MPAs. Total no. of species n = 179.
| Area type | No. of species with<br>at least 33% overlap | No. of species with<br>at least 50 %<br>overlap |
| --- | --- | --- |
| OWF | 0 | 0 |
| MPA | 123 | 62 |
| OWF in MPA | 0 | 0 |
| OWF + MPA | 172 | 103 |
| Uncovered | 139 | 55 |

The biomass-CA of commercial species and P_occ_-CA of sensitive species (i.e., species of conservation concern) were overlaid with OWF and MPA shapes to visualize the potential for co-use (Figure S8) and conservation around OWF (Figure S9).

## Results

### Model performance

The SDM models for P_occ_ had an average AUC of 0.850 (± 0.961 S.D.), an average TSS of 0.626 (± 0.1184 S.D.) and an average MAE of 0.227 (± 0.121 S.D.) (Table S1, Figure S5). The R² from the regression between modelled biomass per km² and biomass values from the validation data was on average 0.112 (± 0.110 S.D.). Thus, the P_occ_-models had an overall better performance than the biomass models.

### Diversity hotspots

Hotspots based on relative P_occ_-CA of demersal fish, epibenthic invertebrates, species of conservation concern (i.e. sensitive species) and commercial species were mostly found along the English, the southern Dutch and the Belgian coast as well as in the deeper, northern parts of the Dutch and German exclusive economic zone (EEZ) (Figure 2). For demersal fish and commercial species, hotspots based on absolute P_occ_-CA were found in the central SNS (Figure 2B&J), for epibenthic invertebrates absolute P_occ_-CA hotspots were found in the Wadden Sea (Figure 2E). Absolute P_occ_-CA (areas with P_occ_ > 0.8) were identified for only eight species (shore crab *Carcinus maenas*, brown shrimp *Crangon crangon*, grey gurnard *Eutrgila gurnardus*, dab *Limanda limanda*, flying crab *Liocarcinus holsatus*, whiting *Merlangius merlangus*, lemon sole *Microstomus kitt* and plaice *Pleuronectes platessa*), and none of these species was a species of conservation concern. Relative P_occ_-CA of species with conservation concern were mostly found in the south-western parts of the SNS, being almost absent from the south-eastern SNS (Figure 2G). Relative biomass-CA were similarly distributed as the relative P_occ_-CA (Figures 2C, F, H, L, but note that in Figure 2F for epibenthic invertebrates only four SDMs were calculated).

**Figure 2.**
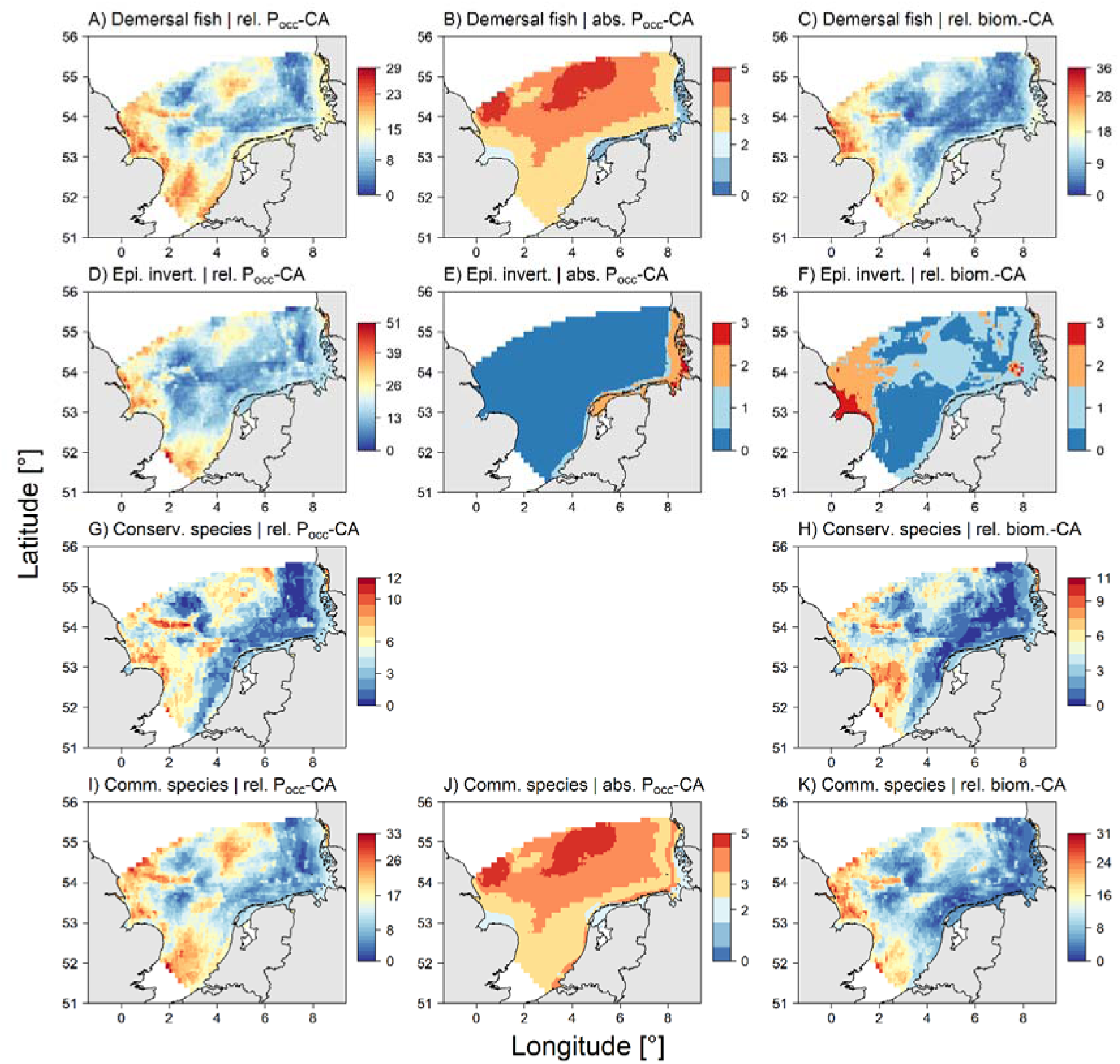
Overlap of species’ core areas to identify hotspots of demersal fish, epibenthic invertebrates, conservation and commercial species diversity. P _occ_-hotspots are based on relative (A, D, G, I) and absolute (B, E, J) core areas. Note that for biomass [kg km ^−2^] (C, F, I, L) absolute core areas could be not calculated and that none of the species of conservation concern had an absolute POC core area (hence the according plot is missing). Grey lines indicate border of national exclusive economic zones.

### Overlap of diversity and OWF & MPA

The overlaps between the distribution of relative P_occ_ hotspots and MPAs indicate that in the British EEZ many hotspots fall within MPA boundaries, whereas in the Dutch and the German EEZ, MPAs overlap only to small extents with hotspots (Figure 3A, C, E & G). Especially relative P_occ_-CA hotspots for sensitive species do not overlap with MPAs in the Dutch and German EEZ (Figure 3E).

**Figure 3.**
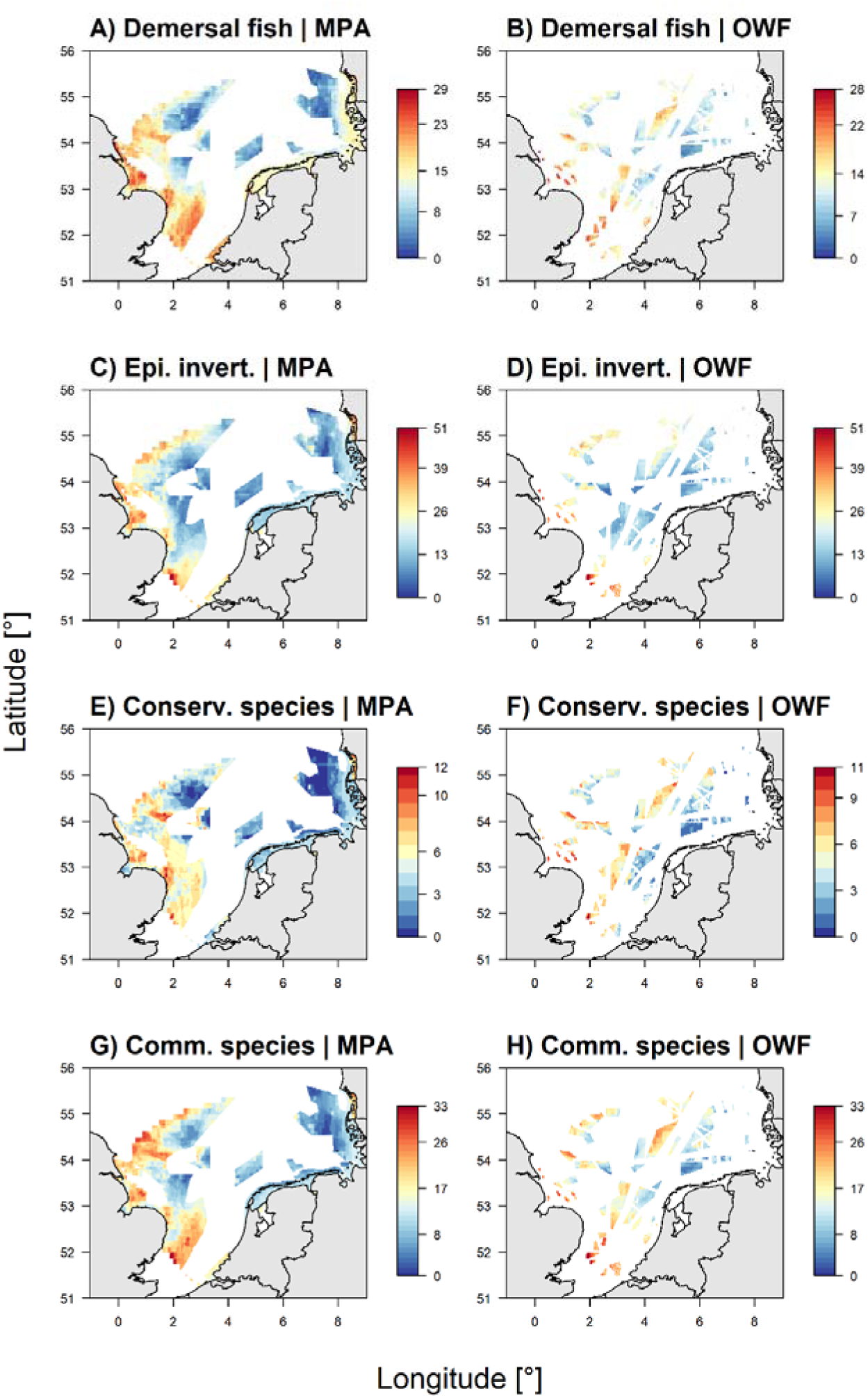
Overlap between relative P _occ_-core areas, designated marine protected areas (MPA) and offshore wind farms (OWF) for demersal fish (A &B), epibenthic invertebrates (C & D), sensitive (E & F) and commercial species (G & H).

Current and future OWF licence areas overlap with several relative P_occ_-hotspots throughout the SNS and could fill gaps in the MPA network coverage (Figure 3 B, D, F &H).

### Single species core areas and overlaps with OWF & MPA

MPA and OWF can be overlaid with single species’ CA to calculate the extent and proportions of overlaps. For example, the relative POC-CA of thornback ray *Raja clavata* had a 20 % overlap with OWFs, 21 % overlap with MPA, 33 % overlap with areas that are designated as both OWF and MPA, whereas 27 % of the thornback rays CA are not covered by neither MPA or OWF (Figure 4).

**Figure 4.**
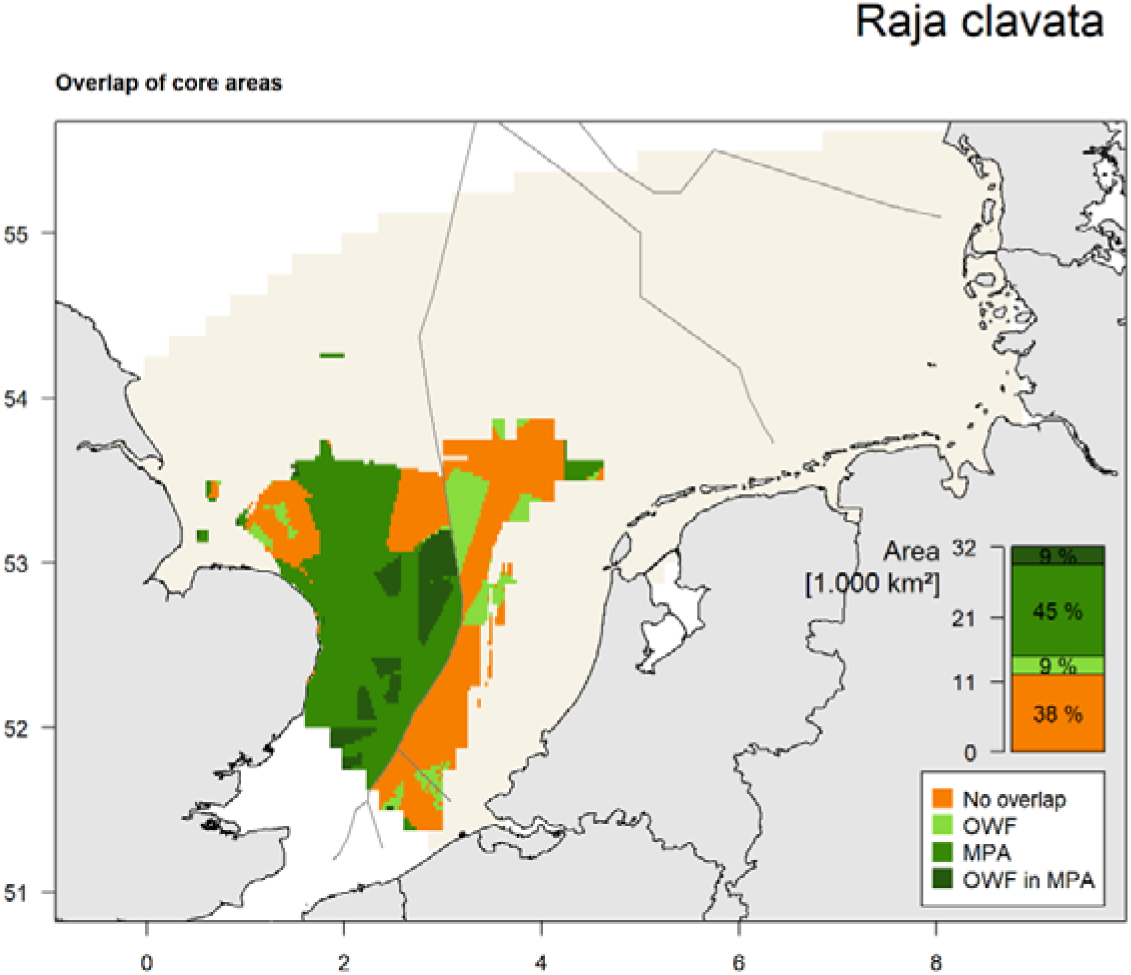
Overlap between the relative core area based on probability of occurrence (P _occ_-CA) of thornback ray Raja clavata and offshore wind parks (OWF) and/or marine protected areas (MPA). In the header the size range of observed thornback rays is indicated (11 cm to 100 cm). The bar plot inset shows total extent in km² of the relative POC-CA as well as percentages of overlap. Blue shade indicates range of study area.

More than half of the species’ P_occ_-CA had an overlap of at least 33 % overlap with MPAs, but 55 species had core areas that were not covered by OWF or MPAs to at least 50% (Table 2). Combining OWF and MPA could lead to a P_occ_-CA of at least 33 % for 172 species accounting for 96 % of all analysed species (n = 179).

### Cluster analysis

The analysis of the scree plot for the cluster analysis indicated that the demersal fish community of the SNS can be portioned into six distinct clusters (Figure S4). These clusters cover the northern English coast (cluster #1), the Wadden Sea (cluster #2), the central northern SNS (cluster #3), the outer and inner coastal waters of Denmark, Germany and the Netherlands (clusters #4 & #6) and the southern English coast (cluster #5) (Figure 5). Of these six clusters, cluster #3, #4 and #6 have only a limited coverage by MPAs, but in each of these clusters OWF are designated.

**Figure 5.**
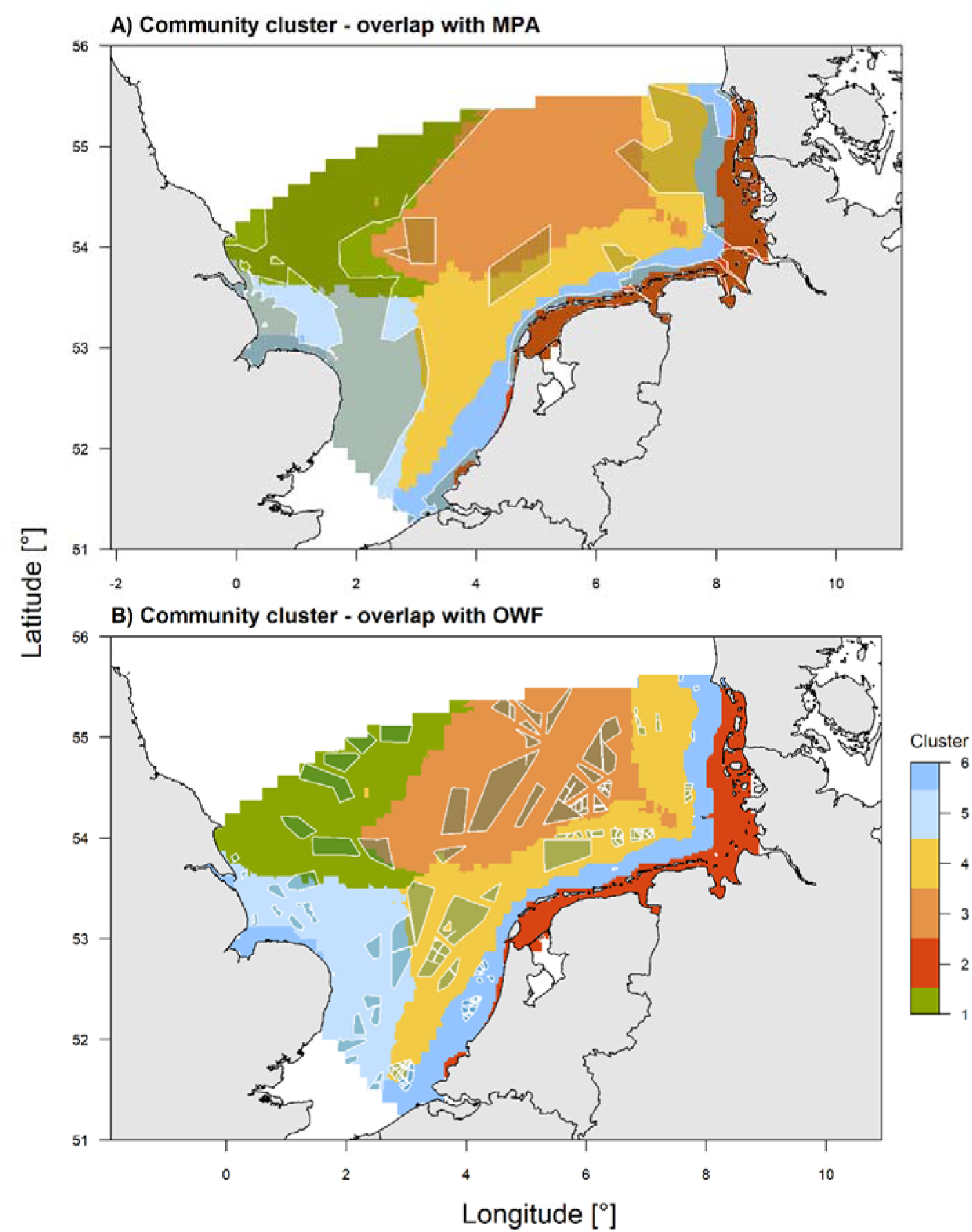
Clusters of the demersal fish community in the southern North Sea. Clusters of similar species are indicated by colours and overlaps with A) marine protected areas (MPA) and B) offshore wind farms (OWF) are shown in darkened shades.

### Responses to single predictors (partial dependencies)

The relationships of P_occ_ vs. the intensity of otter trawling, the intensity of beam trawling, the distance to OWF, bottom temperature and bottom temperature range were analysed for all modelled species (n=179, Figure 6). Further, the responses of 63 demersal fish species are shown in Figure S6.

**Figure 6.**
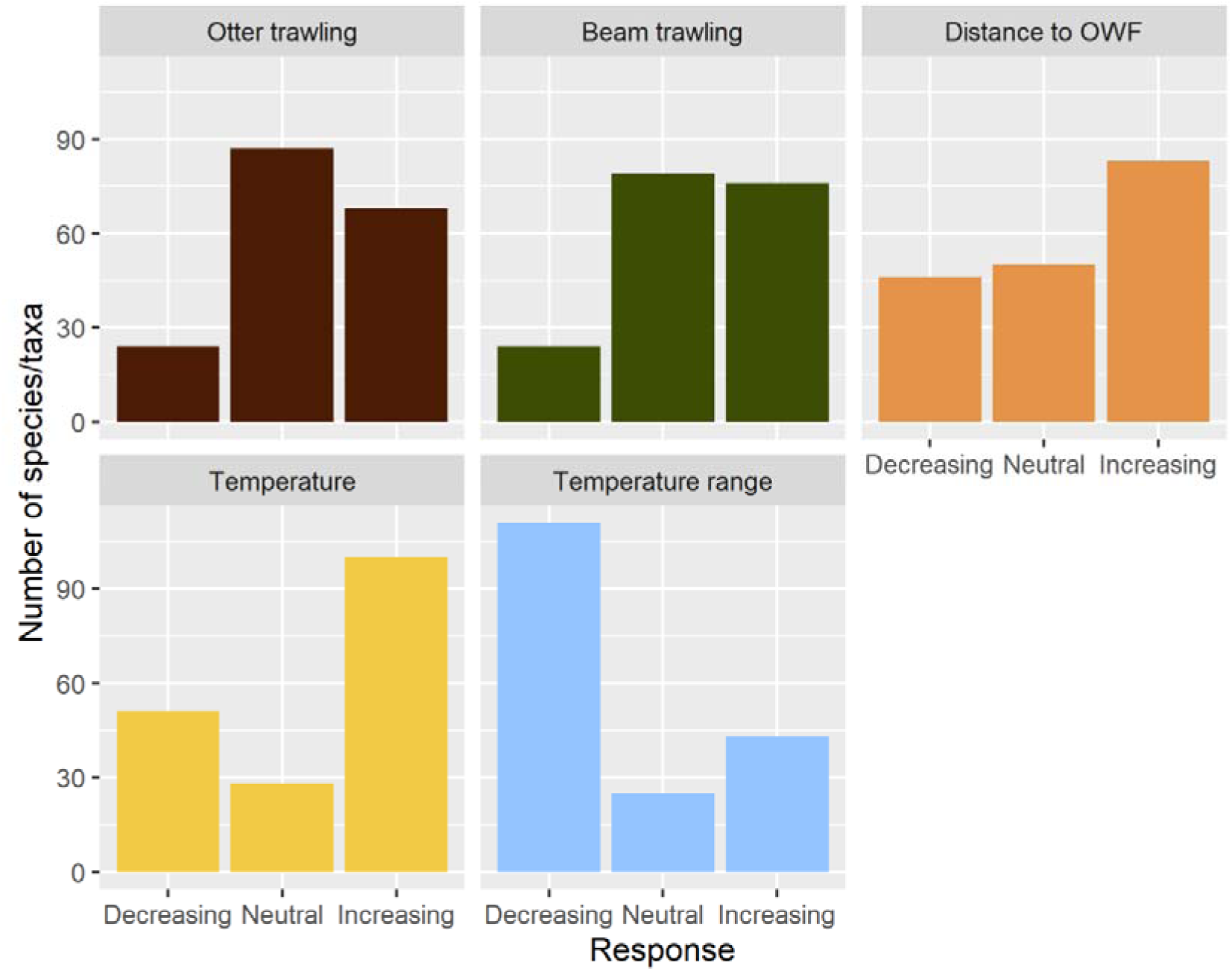
Overview of species’ responses of the probability of occurrence to different predictor variables. OWF = offshore wind farms. Shown are the number of species (n = 179) that show an increasing (significantly positive slope), neutral (slope not significantly different from zero) or decreasing (significantly negative slope) relationship to the predictor variable.

The majority of species had either a relationshsip with a significantly negative (decreasing) or positive slop (increasing), rather than a neutral relationship to any of the aforementioned predictors, indicating that any alteration in the spatial constellation of fishing, offshore wind farm location and temperature will affect more than half of the modelled species.

### Future projections with warming temperatures

SDM were used to model the changes in distribution in three different time slices according to RCP 8.5 and the SDM of several species indicated responses to increasing water temperatures, e.g. red gurnard *Chelidonichthys cuculus* showed an increase in P_occ_ and biomass from the current period (2014 – 2023) until 2100 (Figure S7).

Averaging all modelled P_occ_-values by species within each time period and regressing these averages over time allows calculating the slope as an indication of the change rate in average P_occ_ per year for the suite of 63 demersal fish species (Figure 7). 28 species showed negative changes in P_occ_ between the current time period and the time period of 2090 – 2100, of which some are attributed to the boreal fish fauna such as lemon sole *Microstomus kitt*, haddock *Melanogrammus aeglefinus* or cod *Gadus morhua*. 35 species showed positive changes in average P_occ_, e.g. lesser weaver *Echichthys vipera*, flounder *Platichthys flesus* or sea bass *Dicentrarchus labrax*, which are attributed to the southern (i.e. ‘Lusitanian’) fish fauna of the North Sea.

**Figure 7.**
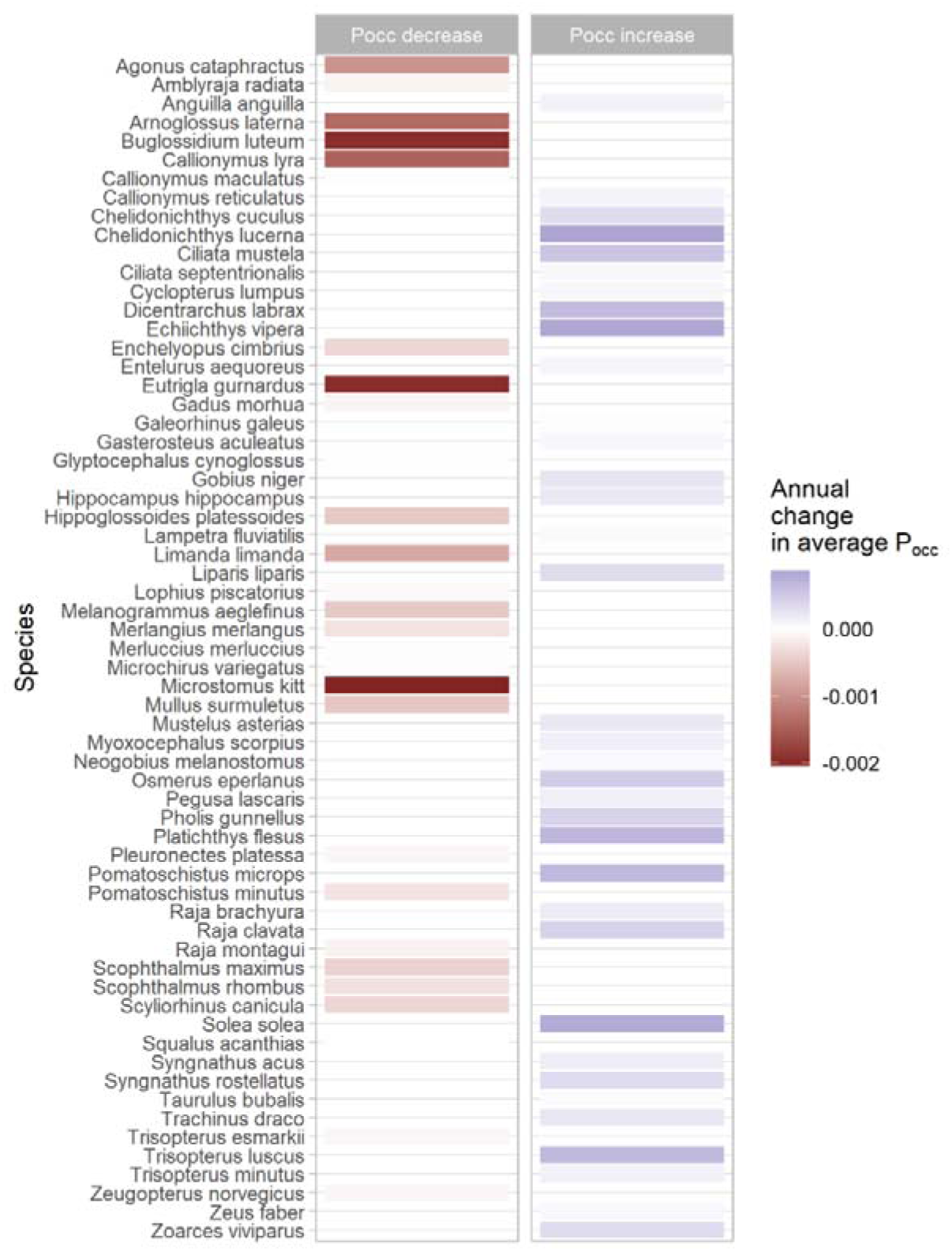
Changes in the annual average probability of occurrence (P _occ_) of 63 modelled demersal fish species in the southern North Sea. Colours indicate the steepness of the slope from a regression between the average P _occ_ and the four modelled time periods (2014 – 2023, 2030 – 2040, 2060 – 2070 and 2090 – 2100).

## Discussion

Our study showed that a considerable proportion species’ core areas and hotspots of diversity overlap with current and future OWF development sites and MPAs, indicating the high potential of enhancing conservation benefits of areas closed to trawled fisheries. However, we show further that many demersal fish species are influenced by human activities and will react to the environmental and socio-economic transformations of the SNS. The majority of taxa responded to gradients in fishing effort, distance to offshore wind farms, and water temperatures. Therefore, the fish and demersal epifauna of the SNS will strongly respond to upcoming changes in any of these aforementioned pressures and actual changes in species’ distributions remain hard to predict as they are subject to more factors than analysed by this study.

In the SNS, fishing is still the human activity with the most widespread spatial extent (Dureuil *et al*. 2018; Couce, Schratzberger & Engelhard 2020; Korpinen *et al*. 2021), but its distribution and intensity will be impacted by the development of OWF (Gusatu *et al*. 2020; Stelzenmüller *et al*. 2022; Bonsu *et al*. 2024) and by the ongoing designation of spatial management measures (i.e. no-take zones) within MPAs (EU 2020; Kriegl *et al*. 2021; Püts *et al*. 2023; UNEP-WCMC & IUCN 2024). However, as shown in a previous study by Probst *et al*. (2021) the currently designated network of MPAs does not cover a substantial proportion of demersal fish species’ core areas and their associated diversity hotspots. Therefore, it is increasingly debated that OWF may contribute to the protection of biodiversity by providing refuge for threatened, rare, sensitive or commercially exploited species such as cod (*Gadus morhua*), rays (*Raja brachyura, R. clavata, R. montagui*), brill *Scophthalmus rhombus*or turbot *Scophthalmus maximus*(Fock 2014; Stelzenmüller *et al*. 2021; Gimpel *et al*. 2023). Our results indicate that several OWF also overlap with the community clusters #3, #4 & #6 and thereby might complement the sparse coverage of MPAs especially in these areas. Our analysis also helps to identify which OWF might offer co-use options for fisheries, e.g. for brown crab *Cancer pagurus*(see also Stelzenmüller *et al*. 2021), lobster *Homarus gammarus* and plaice *Pleuronectes platessa*(Figure S8).

Expected benefits from OWF stem particularly from the artificial reef effect associated with implementing hard structures into habitats dominated by soft sediments and the reduction of mortality through the exclusion of fisheries from OWF areas (Watson *et al*. 2024). However, it has yet to be fully understood if OWFs act only as an attraction for fish biomass or if they can serve as conservation areas with self-sustaining populations (Gimpel *et al*. 2023 and references therein). In addition, benthic communities will likely face potential negative consequences from different pressures associated with the implementation of OWF, such as electro-magnetic changes, noise input or particle motion (Hasselman *et al*. 2023). While there is currently little indication of negative adverse effects of those pressures during the operational phase of OWFs (Svendsen *et al*. 2022), evidence for negative impacts during the commissioning and decommissioning phase suggest mostly adverse impacts on the majority of biota (Li *et al*. 2023; Watson *et al*. 2024).

SDMs allow for deriving species-specific trends of future distribution patterns, which are essential for assessing scenarios of ecosystem states to inform MSP (Stelzenmüller *et al*. 2024). In this study, many demersal fish species showed a response to warming sea bottom temperature, suggesting that the demersal fish community of the SNS might further shift from boreal towards Lusitanian species (Yang 1982; Ehrich & Stransky 2001; Jones *et al*. 2023), even though the turnover of the demersal fish community is already underway. Our results indicate that in the 21^st^ century P will increase for more demersal fish species than decrease, most likely because many boreal species have already retreated from the SNS (Perry *et al*. 2005; Dulvy *et al*. 2008; Engelhard, Righton & Pinnegar 2014). Thereby future projections of boreal species such as cod *Gadus morhua* do not seem to indicate much further change in habitat suitability or P_occ_ (Núñez-Riboni *et al*. 2019, see also Figure 8). In contrast, we predicted that Lusitanian species such as red gurnard (*Chelidonichthys cuculus*), sole (*Solea solea*) or sea bass (*Dicentrarchus labrax*) (Yang 1982) might expand further into the SNS, becoming more common and widespread. But it is important to note that our study only considered the future impact of increasing sea bottom temperature, while the account of changes in other environmental or direct anthropogenic factors may lead to different predictions of species’ distributions.

It is important to note that our study does not attempt to provide a final solution for spatial conservation measures in the southern North Sea, but rather wants to emphasizes the usefulness and essentiality of SDMs for marine spatial conservation planning and that suits of SDMs can be used to derive further data products that may support decisions for MSP. Apart from the here presented analysis, distribution maps from SDMs can also be the foundation of spatial conservation planning tools such as MARXAN (Ball, Possingham & Watts 2009).

Core areas (CA) of species distributions can be defined in various ways, e.g. by resolving seasons (Probst *et al*. 2021). Here we distinguished between relative vs. absolute CA. While absolute CA may be relevant for designating concise areas with species presence known, they come with a drawback: Very rare species, which are often the species of conservation concern, do not occur in such high abundances that allow identifying absolute CAs. Instead, relative CAs can be identified for any species that can be modelled by an SDM. Therefore, for identifying areas of conservation concern, the focus on absolute CA may yield very different and potentially even meaningless results compared to the usage of the relative CA, as many species are rare and endangered.

Our study also analysed CA based on two different metrics i.e. P_occ_ and biomass per km², and demonstrated several advantages of using P_occ_. First, P_occ_ as a target variable allowed us to combine data from multiple surveys, especially from the coastal DYFS, which has not been used by previous studies (Callaway *et al*. 2002; Reiss *et al*. 2010) and allowed us to identify community clusters along the Danish, German and Dutch coast (clusters #1 & #6). Merging data from multiple surveys is often hampered by different gear catchability which can be hard to quantify (Fraser *et al*. 2008), and the use of presence-absence data mitigates the impact of different catchabilities. Secondly, P_occ_ can be applied to more species, as biomass data is not available for many (especially epibenthic) taxa.

Predicted shifts in the distribution ranges of species can have important implications for the assessment of the environmental status as required by the Marine Strategy Framework Directive (MSFD), in which the abundance of many demersal fish species is assessed (Probst, Kloppmann & Kraus 2013; Probst 2023). Setting abundance thresholds for good environmental status (GES) based on historical abundances for species which retreat from an area (Probst & Stelzenmüller 2015; Östman *et al*. 2020) might thereby result in non-achievable status assessment targets. In contrast, for species with increasing abundance, GES-targets based on historical data might lead to overoptimistic assessments. Under changing environmental conditions SDM might therefore allow the identification of more realistic targets for the assessment of the status of ecosystem features rather than analysis based on historical time series. Further, our approach can help to identify ecosystem features, for which the adaptation of assessment targets might be required.

Overall, we show that a comprehensive knowledge on current and future distribution patterns of comprehensive species suites can aid to increase conservation benefits for fish and invertebrates by strategically combining networks of MPAs and OWF, closed for fisheries, in national and cross-border marine spatial planning processes.

## Acknowledgements

This study was funded by the German Federal Ministry of Education and Research as part of the FONA-project “Multiple Stressors on North Sea Life” (MuSSeL, grant numbers 03F0862A, 03F0862D www.mussel-project.de).

## Author contributions

WNP developed research questions, performed modelling and analysis and participated in manuscript writing. VS, JR and CK assisted with data analysis and participated in writing. VS provided resources and funding and participated in writing. HH and HN contributed to data collection and participated in writing. SK, CL and KW provided environmental model data for species distribution modelling and participated in writing.

## Data availability statement

Data on interpolated and modelled species distributions (POC & biomass) in 2014 –2023, 2030 – 2040, 2060 – 70 and 2090 – 2100 are available at Zenodo (https://doi.org/10.5281/zenodo.10069296). Survey data is publicly available at the ICES DATRAS portal: datras.ices.dk.

