## Supplement material for "Current and future relevance of marine spatial planning for the distribution of epibenthic invertebrate and fish species in a heavily used regional sea"

**Supplements content**

**S1** Species list with parameters

**S2** Predictor variables

**S3** Variables for temperature scenarios

**S4** Knee plot for determination of cluster number

**S5** Model performance

**S6** Single species partial responses

**S7** Model output for tub gurnard

**S8** Overlap of commercial taxa and offshore windfarms

**S9** Core areas of species with conservation concern

**Table S1.** List of modelled and summary of evaluation statistics. Also shown are grouping flags for classification as demersal, commercial relevance, conservation concern or neozoa. Parameters a and b refer to the length-weight relationships of the form Weight [kg] = a Length [cm] ^b^. All length-weight-parameters refer to total length in cm except for Cancer pagurus (carapace width in mm), *Homarus gammarus* and *Nephrops norvegicus* (carapace length in mm). Length parameters a & b in bold are from (Wilhelms, 2013). Data on IUCN red list status was accessed via the R-API on 29.04.2024 using the R-package ‘rredlist’ (version 0.7.1). IUCN red list categ*o*ries are NE=’not evaluated’, DD=’data deficient’, LC=’Least concern’, NT=’Near threatened’, VU=’Vulnerable’, EN=’Endangered’, CR=’Critically endangered’. Species were flagged in the column conservation when they were either relevant for national German and OSPAR considerations (see material & methods section) or had an IUCN status = ‘VU’, ‘NT’, ‘EN’ or ‘CR’.

| Species | n | Class | a | b |  | IUCN status | Demersal | Conservation | Commercial | Neozoa | L_min_ [cm] | L_max_ [cm] | AUC | | TSS | MAE | R² biomass vd |
| --- | --- | --- | --- | --- | --- | --- | --- | --- | --- | --- | --- | --- | --- | --- | --- | --- | --- |
| Abra alba | 100 | Bivalvia |  |  |  | NE | 1 | 0 | 0 | 0 | no | no | | 0.758 | 0.430 | 0.290 |  |
| Acanthocardia echinata | 239 | Bivalvia |  |  |  | NE | 1 | 0 | 0 | 0 | no | no | | 0.906 | 0.739 | 0.206 |  |
| Aequipecten opercularis | 274 | Bivalvia |  |  |  | NE | 1 | 0 | 0 | 0 | no | no | | 0.905 | 0.702 | 0.192 |  |
| Aequorea vitrina | 23 | Hydrozoa |  |  |  | NE | 0 | 0 | 0 | 0 | no | no | | 0.957 | 0.922 | 0.078 |  |
| Agonus cataphractus | 3353 | Teleostei | 0.01227 | 2.7953 |  | LC | 1 | 0 | 0 | 0 | 3 | 21 | | 0.693 | 0.270 | 0.377 | 0.155 |
| Alcyonidium diaphanum | 112 | Gymnolaemata |  |  |  | NE | 1 | 0 | 0 | 0 | no | no | | 0.910 | 0.794 | 0.168 |  |
| Alcyonium digitatum | 275 | Anthozoa |  |  |  | NE | 1 | 0 | 0 | 0 | no | no | | 0.890 | 0.639 | 0.294 |  |
| Alloteuthis subulata | 2295 | Cephalopoda | **0.1426** | **1.8222** |  | DD | 0 | 0 | 0 | 0 | 0.5 | 19.5 | | 0.782 | 0.483 | 0.265 | 0.070 |
| Alosa fallax | 129 | Teleostei | 0.00636 | 3.06022 |  | LC | 0 | 1 | 0 | 0 | 5 | 45 | | 0.432 | 0.083 | 0.063 | 0.001 |
| Amblyraja radiata | 93 | Elasmobranchii | 0.01304 | 2.93174 |  | VU | 1 | 1 | 1 | 0 | 11 | 55 | | 0.917 | 0.765 | 0.139 | 0.135 |
| Ammodytes marinus | 209 | Teleostei | 0.00138 | 3.3255 |  | LC | 0 | 0 | 1 | 0 | 5 | 21 | | 0.712 | 0.450 | 0.156 | 0.030 |
| Ammodytes tobianus | 304 | Teleostei | 0.00319 | 2.92034 |  | DD | 0 | 0 | 1 | 0 | 3.5 | 32 | | 0.752 | 0.345 | 0.427 | 0.054 |
| Anguilla anguilla | 47 | Teleostei | 0.0013 | 3.07782 |  | CR | 1 | 1 | 1 | 0 | 6 | 76.2 | | 0.791 | 0.556 | 0.342 | 0.121 |
| Aphrodita aculeata | 594 | Polychaeta |  |  |  | NE | 1 | 0 | 0 | 0 | no | no | | 0.847 | 0.554 | 0.383 |  |
| Arctica islandica | 84 | Bivalvia |  |  |  | NE | 1 | 1 | 0 | 0 | no | no | | 0.942 | 0.824 | 0.174 |  |
| Arnoglossus laterna | 2731 | Teleostei | 0.0063 | 3.07884 |  | LC | 1 | 0 | 0 | 0 | 1 | 35 | | 0.832 | 0.537 | 0.270 | 0.176 |
| Ascidiella aspersa | 21 | Ascidiacea |  |  |  | NE | 1 | 0 | 0 | 0 | no | no | | 0.866 | 0.668 | 0.132 |  |
| Ascidiella scabra | 44 | Ascidiacea |  |  |  | NE | 1 | 0 | 0 | 0 | no | no | | 0.904 | 0.720 | 0.189 |  |
| Asterias rubens | 3843 | Asteroidea |  |  |  | NE | 1 | 0 | 0 | 0 | no | no | | 0.656 | 0.269 | 0.366 |  |
| Astropecten irregularis | 1067 | Asteroidea |  |  |  | NE | 1 | 0 | 0 | 0 | no | no | | 0.877 | 0.640 | 0.243 |  |
| Atelecyclus rotundatus | 28 | Malacostraca |  |  |  | NE | 1 | 0 | 0 | 0 | no | no | | 0.985 | 0.957 | 0.042 |  |
| Atherina presbyter | 22 | Teleostei | 0.00526 | 3.11 |  | LC | 0 | 0 | 0 | 0 | no | no | | 0.772 | 0.504 | 0.329 |  |
| Aurelia aurita | 227 | Scyphozoa |  |  |  | NE | 0 | 0 | 0 | 0 | no | no | | 0.778 | 0.484 | 0.501 |  |
| Belone belone | 76 | Teleostei | 0.00107 | 3.01922 |  | LC | 0 | 0 | 1 | 0 | 7 | 71 | | 0.801 | 0.527 | 0.408 | 0.001 |
| Buccinum undatum | 408 | Gastropoda |  |  |  | NE | 1 | 0 | 1 | 0 | no | no | | 0.874 | 0.631 | 0.156 |  |
| Buglossidium luteum | 2479 | Teleostei | 0.00883 | 3.06289 |  | LC | 1 | 0 | 0 | 0 | 2 | 14 | | 0.903 | 0.690 | 0.182 | 0.182 |
| Calliactis palliata | 26 | Anthozoa |  |  |  | NE | 1 | 0 | 0 | 0 | no | no | | 0.974 | 0.895 | 0.105 |  |
| Callionymus lyra | 3043 | Teleostei | 0.01895 | 2.58876 |  | LC | 1 | 0 | 0 | 0 | 2 | 28 | | 0.836 | 0.572 | 0.228 | 0.117 |
| Callionymus maculatus | 62 | Teleostei | 0.01408 | 2.6554 |  | LC | 1 | 0 | 0 | 0 | 1 | 20 | | 0.836 | 0.575 | 0.421 | 0.008 |
| Callionymus reticulatus | 615 | Teleostei | 0.01742 | 2.51305 |  | LC | 1 | 0 | 0 | 0 | 2.5 | 17.5 | | 0.774 | 0.436 | 0.216 | 0.104 |
| Cancer pagurus | 1774 | Malacostraca | **0.00025** | **2.90285** |  | NE | 1 | 0 | 1 | 0 | 0.5 | 27.5 | | 0.849 | 0.549 | 0.265 | 0.245 |
| Carcinus maenas | 3285 | Malacostraca |  |  |  | NE | 1 | 0 | 0 | 0 | no | no | | 0.937 | 0.715 | 0.144 |  |
| Cerastoderma edule | 196 | Bivalvia |  |  |  | NE | 1 | 0 | 1 | 0 | no | no | | 0.840 | 0.605 | 0.230 |  |
| Chamelea gallina | 23 | Bivalvia |  |  |  | NE | 1 | 0 | 1 | 0 | no | no | | 0.784 | 0.635 | 0.364 |  |
| Chamelea striatula | 127 | Bivalvia |  |  |  | NE | 1 | 0 | 1 | 0 | no | no | | 0.853 | 0.637 | 0.266 |  |
| Chelidonichthys cuculus | 55 | Teleostei | 0.00759 | 3.07974 |  | LC | 1 | 0 | 1 | 0 | 5 | 31 | | 0.960 | 0.811 | 0.106 | 0.173 |
| Chelidonichthys lucerna | 2154 | Teleostei | 0.0088 | 2.9989 |  | LC | 1 | 0 | 1 | 0 | 1 | 56 | | 0.748 | 0.369 | 0.316 | 0.112 |
| Chrysaora hysoscella | 499 | Scyphozoa |  |  |  | NE | 0 | 0 | 0 | 0 | no | no | | 0.759 | 0.373 | 0.301 |  |
| Ciliata mustela | 1987 | Teleostei | 0.00577 | 3.0845 |  | LC | 1 | 0 | 0 | 0 | 2 | 25 | | 0.831 | 0.510 | 0.332 | 0.233 |
| Ciliata septentrionalis | 24 | Teleostei | 0.00827 | 3.1785 |  | LC | 1 | 0 | 0 | 0 | 7 | 21 | | 0.745 | 0.609 | 0.391 | 0.029 |
| Ciona intestinalis | 63 | Ascidiacea |  |  |  | NE | 1 | 0 | 0 | 0 | no | no | | 0.871 | 0.648 | 0.104 |  |
| Clupea harengus | 4235 | Teleostei | 0.00632 | 3.07354 |  | LC | 0 | 0 | 1 | 0 | 1 | 35 | | 0.707 | 0.295 | 0.345 | 0.023 |
| Corystes cassivelaunus | 778 | Malacostraca |  |  |  | NE | 1 | 0 | 0 | 0 | no | no | | 0.876 | 0.632 | 0.252 |  |
| Crangon allmanni | 133 | Malacostraca |  |  |  | NE | 1 | 0 | 1 | 0 | no | no | | 0.810 | 0.485 | 0.447 |  |
| Crangon crangon | 4549 | Malacostraca | **0.0045** | **3.257** |  | NE | 1 | 0 | 1 | 0 | 0.4 | 12.6 | | 0.958 | 0.841 | 0.072 | 0.532 |
| Crepidula fornicata | 252 | Gastropoda |  |  |  | NE | 1 | 0 | 0 | 0 | no | no | | 0.800 | 0.512 | 0.264 |  |
| Crossaster papposus | 60 | Asteroidea |  |  |  | NE | 1 | 0 | 0 | 0 | no | no | | 0.985 | 0.909 | 0.090 |  |
| Cyanea capillata | 59 | Scyphozoa |  |  |  | NE | 0 | 0 | 0 | 0 | no | no | | 0.839 | 0.651 | 0.346 |  |
| Cyanea lamarckii | 106 | Scyphozoa |  |  |  | NE | 0 | 0 | 0 | 0 | no | no | | 0.852 | 0.607 | 0.276 |  |
| Cyclopterus lumpus | 40 | Teleostei | 0.07978 | 2.7979 |  | NT | 1 | 1 | 0 | 0 | 3 | 46 | | 0.448 | 0.190 | 0.806 | 0.000 |
| Cylista troglodytes | 36 | Anthozoa |  |  |  | NE | 1 | 0 | 0 | 0 | no | no | | 0.781 | 0.555 | 0.223 |  |
| Dicentrarchus labrax | 143 | Teleostei | 0.01093 | 3.01067 |  | LC | 1 | 0 | 1 | 0 | 4 | 75 | | 0.852 | 0.544 | 0.164 | 0.075 |
| Diogenes pugilator | 464 | Malacostraca |  |  |  | NE | 1 | 0 | 0 | 1 | no | no | | 0.866 | 0.593 | 0.277 |  |
| Donax vittatus | 208 | Bivalvia |  |  |  | NE | 1 | 0 | 0 | 0 | no | no | | 0.901 | 0.688 | 0.286 |  |
| Dosinia exoleta | 29 | Bivalvia |  |  |  | NE | 1 | 0 | 0 | 0 | no | no | | 0.878 | 0.747 | 0.252 |  |
| Echiichthys vipera | 1417 | Teleostei | 0.01317 | 2.99728 |  | LC | 1 | 0 | 0 | 0 | 1 | 88 | | 0.912 | 0.679 | 0.177 | 0.227 |
| Echinocardium cordatum | 730 | Echinoidea |  |  |  | NE | 1 | 0 | 0 | 0 | no | no | | 0.746 | 0.392 | 0.465 |  |
| Echinus esculentus | 22 | Echinoidea |  |  |  | N | 1 | 0 | 0 | 0 | no | no | | 0.959 | 0.819 | 0.015 |  |
| Eledone cirrhosa | 27 | Cephalopoda | **0.6971** | **2.4196** |  | LC | 1 | 0 | 0 | 0 | no | no | | 0.904 | 0.773 | 0.226 |  |
| Enchelyopus cimbrius | 569 | Teleostei | 0.00368 | 3.0842 |  | LC | 1 | 0 | 0 | 0 | 7 | 29 | | 0.944 | 0.760 | 0.167 | 0.184 |
| Engraulis encrasicolus | 494 | Teleostei | 0.00549 | 3.05873 |  | LC | 0 | 0 | 1 | 0 | 2 | 20 | | 0.813 | 0.577 | 0.293 | 0.005 |
| Ensis leei | 37 | Bivalvia |  |  |  | NE | 1 | 0 | 1 | 0 | no | no | | 0.629 | 0.429 | 0.128 |  |
| Ensis siliqua | 25 | Bivalvia |  |  |  | NE | 1 | 0 | 1 | 0 | no | no | | 0.650 | 0.382 | 0.616 |  |
| Entelurus aequoreus | 65 | Teleostei | 0.00026 | 2.89035 |  | LC | 1 | 0 | 0 | 0 | 7 | 50 | | 0.817 | 0.512 | 0.273 | 0.052 |
| Euspira catena | 138 | Gastropoda |  |  |  | NE | 1 | 0 | 0 | 0 | no | no | | 0.877 | 0.596 | 0.369 |  |
| Euspira nitida | 20 | Gastropoda |  |  |  | NE | 1 | 0 | 0 | 0 | no | no | | 0.676 | 0.384 | 0.216 |  |
| Eutrigla gurnardus | 2297 | Teleostei | 0.00725 | 3.05588 |  | LC | 1 | 0 | 1 | 0 | 4 | 44 | | 0.965 | 0.858 | 0.069 | 0.293 |
| Flustra foliacea | 105 | Gymnolaemata |  |  |  | NE | 1 | 0 | 0 | 0 | no | no | | 0.934 | 0.767 | 0.155 |  |
| Gadus morhua | 1007 | Teleostei | 0.00714 | 3.07736 |  | LC | 1 | 1 | 1 | 0 | 5 | 113 | | 0.713 | 0.336 | 0.294 | 0.029 |
| Galeorhinus galeus | 55 | Elasmobranchii | 0.00618 | 2.95169 |  | CR | 1 | 1 | 1 | 0 | 30 | 168 | | 0.948 | 0.830 | 0.169 | 0.050 |
| Gasterosteus aculeatus | 157 | Teleostei | 0.01008 | 3.19168 |  | LC | 1 | 0 | 0 | 0 | 0.3 | 7.6 | | 0.793 | 0.550 | 0.285 | 0.019 |
| Glyptocephalus cynoglossus | 32 | Teleostei | 0.00218 | 3.29828 |  | VU | 1 | 1 | 1 | 0 | 5 | 35 | | 0.969 | 0.915 | 0.084 | 0.006 |
| Gobius niger | 27 | Teleostei | 0.01138 | 3.05172 |  | LC | 1 | 0 | 0 | 0 | 6 | 13.5 | | 0.896 | 0.768 | 0.032 | 0.188 |
| Goneplax rhomboides | 208 | Malacostraca |  |  |  | NE | 1 | 0 | 0 | 0 | no | no | | 0.948 | 0.852 | 0.108 |  |
| Hemigrapsus takanoi | 20 | Malacostraca |  |  |  | NE | 1 | 0 | 0 | 0 | no | no | | 0.938 | 0.872 | 0.128 |  |
| Henricia sanguinolenta | 36 | Asteroidea |  |  |  | NE | 1 | 0 | 0 | 0 | no | no | | 0.995 | 0.991 | 0.009 |  |
| Hippocampus hippocampus | 55 | Teleostei | 0.0024 | 3.0015 |  | DD | 1 | 1 | 0 | 0 | no | no | | 0.894 | 0.671 | 0.186 |  |
| Hippoglossoides platessoides | 372 | Teleostei | 0.00299 | 3.25901 |  | EN | 1 | 1 | 1 | 0 | 4 | 32 | | 0.973 | 0.863 | 0.091 | 0.219 |
| Homarus gammarus | 220 | Malacostraca | 0.00131 | 2.882 |  | LC | 1 | 1 | 1 | 0 | 2.3 | 20 | | 0.931 | 0.714 | 0.189 | 0.293 |
| Hyas araneus | 85 | Malacostraca |  |  |  | NE | 1 | 0 | 0 | 0 | no | no | | 0.908 | 0.739 | 0.212 |  |
| Hyas coarctatus | 82 | Malacostraca |  |  |  | NE | 1 | 0 | 0 | 0 | no | no | | 0.958 | 0.871 | 0.128 |  |
| Hydractinia echinata | 80 | Hydrozoa |  |  |  | NE | 1 | 0 | 0 | 0 | no | no | | 0.759 | 0.464 | 0.139 |  |
| Hyperoplus lanceolatus | 1248 | Teleostei | 0.00474 | 2.81235 |  | LC | 0 | 0 | 1 | 0 | 4 | 37 | | 0.784 | 0.435 | 0.371 | 0.023 |
| Illex coindetii | 104 | Cephalopoda | **0.0495** | **2.9526** |  | LC | 0 | 0 | 1 | 0 | 2.5 | 18.5 | | 0.912 | 0.734 | 0.223 | 0.045 |
| Inachus dorsettensis | 72 | Malacostraca |  |  |  | NE | 1 | 0 | 0 | 0 | no | no | | 0.965 | 0.844 | 0.100 |  |
| Laevicardium crassum | 44 | Bivalvia |  |  |  | NE | 1 | 0 | 0 | 0 | no | no | | 0.788 | 0.514 | 0.483 |  |
| Lampetra fluviatilis | 100 | Petromyzonti | 0.00102 | 3.1659 |  | LC | 1 | 1 | 0 | 0 | 15 | 40 | | 0.765 | 0.521 | 0.295 | 0.005 |
| Limanda limanda | 5665 | Teleostei | 0.00713 | 3.11283 |  | LC | 1 | 0 | 1 | 0 | 1 | 37 | | 0.910 | 0.729 | 0.151 | 0.098 |
| Liocarcinus depurator | 1010 | Malacostraca |  |  |  | NE | 1 | 0 | 0 | 0 | no | no | | 0.823 | 0.571 | 0.294 |  |
| Liocarcinus holsatus | 4877 | Malacostraca |  |  |  | NE | 1 | 0 | 0 | 0 | no | no | | 0.785 | 0.483 | 0.293 |  |
| Liocarcinus marmoreus | 173 | Malacostraca |  |  |  | NE | 1 | 0 | 0 | 0 | no | no | | 0.786 | 0.536 | 0.298 |  |
| Liocarcinus navigator | 354 | Malacostraca |  |  |  | NE | 1 | 0 | 0 | 0 | no | no | | 0.884 | 0.686 | 0.217 |  |
| Liocarcinus vernalis | 78 | Malacostraca |  |  |  | NE | 1 | 0 | 0 | 0 | no | no | | 0.836 | 0.540 | 0.259 |  |
| Liparis liparis | 760 | Teleostei | 0.03016 | 3 |  | LC | 1 | 0 | 0 | 0 | 2 | 18 | | 0.783 | 0.435 | 0.466 | 0.074 |
| Loligo forbesii | 490 | Cephalopoda | **0.1122** | **2.5261** |  | LC | 0 | 0 | 1 | 0 | 1 | 38 | | 0.882 | 0.662 | 0.280 | 0.077 |
| Loligo vulgaris | 281 | Cephalopoda | **0.1961** | **2.3632** |  | DD | 0 | 0 | 1 | 0 | 2 | 40 | | 0.818 | 0.589 | 0.353 | 0.006 |
| Lophius piscatorius | 73 | Teleostei | 0.0142 | 2.91911 |  | LC | 1 | 0 | 1 | 0 | 14 | 59 | | 0.946 | 0.823 | 0.175 | 0.067 |
| Luidia sarsii | 298 | Asteroidea |  |  |  | NE | 1 | 0 | 0 | 0 | no | no | | 0.950 | 0.822 | 0.146 |  |
| Lutraria lutraria | 63 | Bivalvia |  |  |  | NE | 1 | 0 | 0 | 0 | no | no | | 0.779 | 0.496 | 0.192 |  |
| Macoma balthica | 243 | Bivalvia |  |  |  | NE | 1 | 0 | 0 | 0 | no | no | | 0.835 | 0.537 | 0.248 |  |
| Macropodia rostrata | 439 | Malacostraca |  |  |  | NE | 1 | 0 | 0 | 0 | no | no | | 0.794 | 0.440 | 0.375 |  |
| Macropodia tenuirostris | 62 | Malacostraca |  |  |  | NE | 1 | 0 | 0 | 0 | no | no | | 0.837 | 0.578 | 0.173 |  |
| Mactra corallina | 20 | Bivalvia |  |  |  | NE | 1 | 0 | 0 | 0 | no | no | | 0.981 | 0.923 | 0.076 |  |
| Mactra stultorum | 229 | Bivalvia |  |  |  | NE | 1 | 0 | 0 | 0 | no | no | | 0.800 | 0.513 | 0.374 |  |
| Magallana gigas | 113 | Bivalvia |  |  |  | NE | 1 | 0 | 1 | 1 | no | no | | 0.923 | 0.736 | 0.191 |  |
| Maja squinado | 36 | Malacostraca |  |  |  | NE | 1 | 0 | 0 | 0 | no | no | | 0.994 | 0.991 | 0.009 |  |
| Melanogrammus aeglefinus | 415 | Teleostei | 0.00583 | 3.13252 |  | LC | 1 | 0 | 1 | 0 | 8 | 47 | | 0.924 | 0.726 | 0.172 | 0.145 |
| Merlangius merlangus | 5567 | Teleostei | 0.00707 | 3.04549 |  | LC | 1 | 0 | 1 | 0 | 1 | 51 | | 0.739 | 0.356 | 0.273 | 0.038 |
| Merluccius merluccius | 27 | Teleostei | 0.00517 | 3.10575 |  | LC | 1 | 0 | 1 | 0 | 39 | 74 | | 0.928 | 0.883 | 0.117 | 0.027 |
| Metridium dianthus | 34 | Anthozoa |  |  |  | NE | 1 | 0 | 0 | 0 | no | no | | 0.910 | 0.734 | 0.265 |  |
| Microchirus variegatus | 67 | Teleostei | 0.00825 | 3.09361 |  | LC | 1 | 0 | 0 | 0 | 6.5 | 19 | | 0.979 | 0.905 | 0.094 | 0.190 |
| Micromesistius poutassou | 37 | Teleostei | 0.00447 | 3.10521 |  | LC | 0 | 0 | 1 | 0 | 11 | 26 | | 0.770 | 0.558 | 0.440 | 0.071 |
| Microstomus kitt | 1837 | Teleostei | 0.0104 | 3.02375 |  | LC | 1 | 0 | 1 | 0 | 5 | 37 | | 0.913 | 0.714 | 0.129 | 0.410 |
| Mnemiopsis leidyi | 74 | Tentaculata |  |  |  | NE | 0 | 0 | 0 | 1 | no | no | | 0.800 | 0.649 | 0.239 |  |
| Molgula occulta | 48 | Ascidiacea |  |  |  | NE | 1 | 0 | 0 | 0 | no | no | | 0.969 | 0.916 | 0.084 |  |
| Mullus surmuletus | 1014 | Teleostei | 0.00782 | 3.16344 |  | LC | 1 | 0 | 1 | 0 | 4 | 31 | | 0.827 | 0.537 | 0.215 | 0.051 |
| Mustelus asterias | 269 | Elasmobranchii | 0.00148 | 3.19721 |  | NT | 1 | 1 | 1 | 0 | 23 | *125* | | 0.*917* | 0.*703* | 0.*230* | 0.0*33* |
| Myoxocephalus scorpius | 1979 | Teleostei | 0.00967 | 3.15909 |  | LC | 1 | 0 | 0 | 0 | 1 | 29 | | 0.698 | 0.326 | 0.309 | 0.096 |
| Mytilus edulis | 473 | Bivalvia |  |  |  | NE | 1 | 0 | 1 | 0 | no | no | | 0.807 | 0.500 | 0.217 |  |
| Necora puber | 443 | Malacostraca |  |  |  | NE | 1 | 0 | 0 | 0 | no | no | | 0.869 | 0.606 | 0.244 |  |
| Neogobius melanostomus | 27 | Teleostei | 0.01324 | 3.04221 |  | LC | 1 | 0 | 0 | 1 | 5 | 17 | | 0.777 | 0.682 | 0.118 | 0.001 |
| Nephrops norvegicus | 348 | Malacostraca | **0.00042** | **3.11965** |  | LC | 1 | 1 | 1 | 0 | 0.4 | 10.2 | | 0.976 | 0.865 | 0.053 | 0.350 |
| Neptunea antiqua | 37 | Gastropoda |  |  |  | NE | 1 | 0 | 1 | 0 | no | no | | 0.990 | 0.970 | 0.030 |  |
| Ophiothrix fragilis | 176 | Ophiuroidea |  |  |  | NE | 1 | 0 | 0 | 0 | no | no | | 0.927 | 0.792 | 0.116 |  |
| Ophiura albida | 634 | Ophiuroidea |  |  |  | NE | 1 | 0 | 0 | 0 | no | no | | 0.807 | 0.509 | 0.313 |  |
| Ophiura ophiura | 2426 | Ophiuroidea |  |  |  | NE | 1 | 0 | 0 | 0 | no | no | | 0.717 | 0.337 | 0.336 |  |
| Osmerus eperlanus | 1754 | Teleostei | 0.00462 | 3.14993 |  | LC | 1 | 0 | 1 | 0 | 1 | 24 | | 0.905 | 0.644 | 0.237 | 0.178 |
| Pagurus bernhardus | 2671 | Malacostraca |  |  |  | NE | 1 | 0 | 0 | 0 | no | no | | 0.671 | 0.260 | 0.354 |  |
| Pagurus longicarpus | 55 | Malacostraca |  |  |  | NE | 1 | 0 | 0 | 1 | no | no | | 0.897 | 0.794 | 0.063 |  |
| Pagurus prideaux | 49 | Malacostraca |  |  |  | NE | 1 | 0 | 0 | 0 | no | no | | 0.961 | 0.883 | 0.033 |  |
| Palaemon macrodactylus | 30 | Malacostraca |  |  |  | NE | 1 | 0 | 1 | 1 | no | no | | 0.963 | 0.851 | 0.149 |  |
| Palaemon serratus | 203 | Malacostraca |  |  |  | NE | 1 | 0 | 1 | 0 | no | no | | 0.784 | 0.440 | 0.304 |  |
| Pandalus montagui | 433 | Malacostraca |  |  |  | NE | 1 | 0 | 0 | 0 | no | no | | 0.775 | 0.438 | 0.293 |  |
| Paraleptopentacta elongata | 34 | Holothuroidea |  |  |  | NE | 1 | 0 | 1 | 0 | no | no | | 0.963 | 0.831 | 0.044 |  |
| Pecten maximus | 70 | Bivalvia |  |  |  | NE | 1 | 0 | 1 | 0 | no | no | | 0.950 | 0.826 | 0.063 |  |
| Pegusa lascaris | 21 | Teleostei | 0.00669 | 3.1488 |  | LC | 1 | 0 | 0 | 0 | 9 | 30 | | 0.861 | 0.793 | 0.207 | 0.000 |
| Pholis gunnellus | 648 | Teleostei | 0.00342 | 3.0575 |  | LC | 1 | 0 | 0 | 0 | 4 | 21 | | 0.789 | 0.450 | 0.299 | 0.091 |
| Pilumnus hirtellus | 43 | Malacostraca |  |  |  | NE | 1 | 0 | 0 | 0 | no | no | | 0.854 | 0.588 | 0.230 |  |
| Pisidia longicornis | 86 | Malacostraca |  |  |  | NE | 1 | 0 | 0 | 0 | no | no | | 0.887 | 0.671 | 0.192 |  |
| Platichthys flesus | 2577 | Teleostei | 0.01004 | 3.02906 |  | LC | 1 | 0 | 1 | 0 | 2 | 43.5 | | 0.760 | 0.402 | 0.310 | 0.152 |
| Pleurobrachia pileus | 91 | Tentaculata |  |  |  | NE | 0 | 0 | 0 | 0 | no | no | | 0.924 | 0.730 | 0.182 |  |
| Pleuronectes platessa | 7169 | Teleostei | 0.00936 | 3.05611 |  | LC | 1 | 0 | 1 | 0 | 1 | 83.5 | | 0.765 | 0.408 | 0.261 | 0.272 |
| Pomatoschistus microps | 73 | Teleostei | 0.0075 | 3.18 |  | LC | 1 | 0 | 0 | 0 | 2 | 7 | | 0.875 | 0.637 | 0.298 | 0.012 |
| Pomatoschistus minutus | 1809 | Teleostei | 0.00662 | 3.06687 |  | LC | 1 | 0 | 0 | 0 | 2 | 10.5 | | 0.921 | 0.776 | 0.134 | 0.206 |
| Portumnus latipes | 72 | Malacostraca |  |  |  | NE | 1 | 0 | 0 | 0 | no | no | | 0.807 | 0.526 | 0.142 |  |
| Psammechinus miliaris | 488 | Echinoidea |  |  |  | NE | 1 | 0 | 0 | 0 | no | no | | 0.797 | 0.491 | 0.241 |  |
| Raja brachyura | 155 | Elasmobranchii | 0.00211 | 3.29 |  | NT | 1 | 1 | 1 | 0 | 9.5 | 100 | | 0.976 | 0.882 | 0.090 | 0.102 |
| Raja clavata | 619 | Elasmobranchii | 0.00204 | 3.24322 |  | NT | 1 | 1 | 1 | 0 | 11 | 100 | | 0.933 | 0.718 | 0.171 | 0.199 |
| Raja montagui | 557 | Elasmobranchii | 0.00218 | 3.31405 |  | LC | 1 | 1 | 1 | 0 | 11 | 82 | | 0.927 | 0.750 | 0.176 | 0.151 |
| Rhizostoma pulmo | 550 | Scyphozoa |  |  |  | NE | 0 | 0 | 0 | 0 | no | no | | 0.850 | 0.573 | 0.297 |  |
| Sardina pilchardus | 221 | Teleostei | 0.00564 | 3.11987 |  | NT | 0 | 1 | 1 | 0 | 4 | 39 | | 0.780 | 0.545 | 0.371 | 0.004 |
| Scomber scombrus | 709 | Teleostei | 0.00357 | 3.25044 |  | LC | 0 | 0 | 1 | 0 | 5 | 44 | | 0.854 | 0.608 | 0.300 | 0.036 |
| Scophthalmus maximus | 1146 | Teleostei | 0.01339 | 3.07153 |  | LC | 1 | 0 | 1 | 0 | 3 | 88 | | 0.777 | 0.419 | 0.282 | 0.098 |
| Scophthalmus rhombus | 619 | Teleostei | 0.01359 | 3.01258 |  | LC | 1 | 0 | 1 | 0 | 3 | 57 | | 0.719 | 0.322 | 0.481 | 0.061 |
| Scyliorhinus canicula | 1064 | Elasmobranchii | 0.00138 | 3.20749 |  | LC | 1 | 1 | 0 | 0 | 9 | 74 | | 0.940 | 0.753 | 0.133 | 0.288 |
| Sepia officinalis | 153 | Cephalopoda |  |  |  | LC | 1 | 0 | 1 | 0 | no | no | | 0.913 | 0.676 | 0.166 |  |
| Sepiola atlantica | 311 | Cephalopoda |  |  |  | DD | 1 | 0 | 0 | 0 | no | no | | 0.672 | 0.244 | 0.290 |  |
| Solea solea | 3300 | Teleostei | 0.00731 | 3.06022 |  | DD | 1 | 0 | 1 | 0 | 2 | 44 | | 0.752 | 0.406 | 0.311 | 0.362 |
| Spatangus purpureus | 42 | Echinoidea |  |  |  | NE | 1 | 0 | 0 | 0 | no | no | | 0.902 | 0.678 | 0.122 |  |
| Spisula elliptica | 24 | Bivalvia |  |  |  | NE | 1 | 0 | 1 | 0 | no | no | | 0.922 | 0.849 | 0.150 |  |
| Spisula solida | 147 | Bivalvia |  |  |  | NE | 1 | 0 | 1 | 0 | no | no | | 0.745 | 0.450 | 0.150 |  |
| Spisula subtruncata | 132 | Bivalvia |  |  |  | NE | 1 | 0 | 1 | 0 | no | no | | 0.809 | 0.516 | 0.154 |  |
| Sprattus sprattus | 2553 | Teleostei | 0.00538 | 3.11107 |  | LC | 0 | 0 | 1 | 0 | 2.5 | 18 | | 0.707 | 0.332 | 0.337 | 0.023 |
| Squalus acanthias | 32 | Elasmobranchii | 0.00273 | 3.08469 |  | VU | 1 | 1 | 1 | 0 | 26 | 121 | | 0.846 | 0.660 | 0.339 | 0.000 |
| Syngnathus acus | 325 | Teleostei | 0.00022 | 3.25084 |  | LC | 1 | 0 | 0 | 0 | 5 | 46 | | 0.830 | 0.528 | 0.183 | 0.155 |
| Syngnathus rostellatus | 2365 | Teleostei | 0.00014 | 3.41123 |  | DD | 1 | 0 | 0 | 0 | 1 | 23 | | 0.830 | 0.524 | 0.315 | 0.144 |
| Taurulus bubalis | 162 | Teleostei | 0.0154 | 3 |  | LC | 1 | 0 | 0 | 0 | 3.5 | 23 | | 0.811 | 0.530 | 0.208 | 0.102 |
| Todaropsis eblanae | 33 | Cephalopoda | **0.1681** | **2.595** |  | LC | 0 | 0 | 1 | 0 | 8 | 17 | | 0.917 | 0.844 | 0.156 | 0.001 |
| Trachinus draco | 76 | Teleostei | 0.0061 | 3.04751 |  | LC | 1 | 0 | 0 | 0 | 8 | 41 | | 0.933 | 0.769 | 0.114 | 0.085 |
| Trachurus trachurus | 1416 | Teleostei | 0.00925 | 2.96653 |  | LC | 0 | 0 | 1 | 0 | 1 | 93 | | 0.681 | 0.282 | 0.354 | 0.039 |
| Trisopterus esmarkii | 87 | Teleostei | 0.00518 | 3.10557 |  | LC | 1 | 0 | 1 | 0 | 6 | 20 | | 0.949 | 0.794 | 0.154 | 0.021 |
| Trisopterus luscus | 823 | Teleostei | 0.00888 | 3.11625 |  | LC | 1 | 0 | 1 | 0 | 3 | 36.5 | | 0.813 | 0.479 | 0.244 | 0.070 |
| Trisopterus minutus | 320 | Teleostei | 0.00696 | 3.13903 |  | LC | 1 | 0 | 1 | 0 | 3 | 24 | | 0.950 | 0.756 | 0.104 | 0.163 |
| Tritia reticulata | 178 | Gastropoda |  |  |  | NE | 1 | 0 | 0 | 0 | no | no | | 0.929 | 0.778 | 0.109 |  |
| Turritellinella tricarinata | 30 | Gastropoda |  |  |  | NE | 1 | 0 | 0 | 0 | no | no | | 0.970 | 0.909 | 0.090 |  |
| Upogebia deltaura | 28 | Malacostraca |  |  |  | NE | 1 | 1 | 0 | 0 | no | no | | 0.976 | 0.954 | 0.046 |  |
| Zeugopterus norvegicus | 67 | Teleostei | 0.01288 | 2.9795 |  | NE | 1 | 0 | 0 | 0 | 6 | 15 | | 0.896 | 0.666 | 0.134 | 0.008 |
| Zeus faber | 54 | Teleostei | 0.02371 | 2.88391 |  | DD | 1 | 0 | 1 | 0 | 10 | 36 | | 0.916 | 0.725 | 0.273 | 0.063 |
| Zoarces viviparus | 721 | Teleostei | 0.00781 | 2.83177 |  | LC | 1 | 0 | 0 | 0 | 1 | 30 | | 0.904 | 0.650 | 0.176 | 0.288 |

Wilhelms, I. 2013. Atlas of length-weight relationships of 93 fish and crustacean species from the North Sea and the North-East Atlantic. Thünen Working Paper: 12. 552 pp.

**Figure S2.** Predictor variables for SDMs. For details see table.

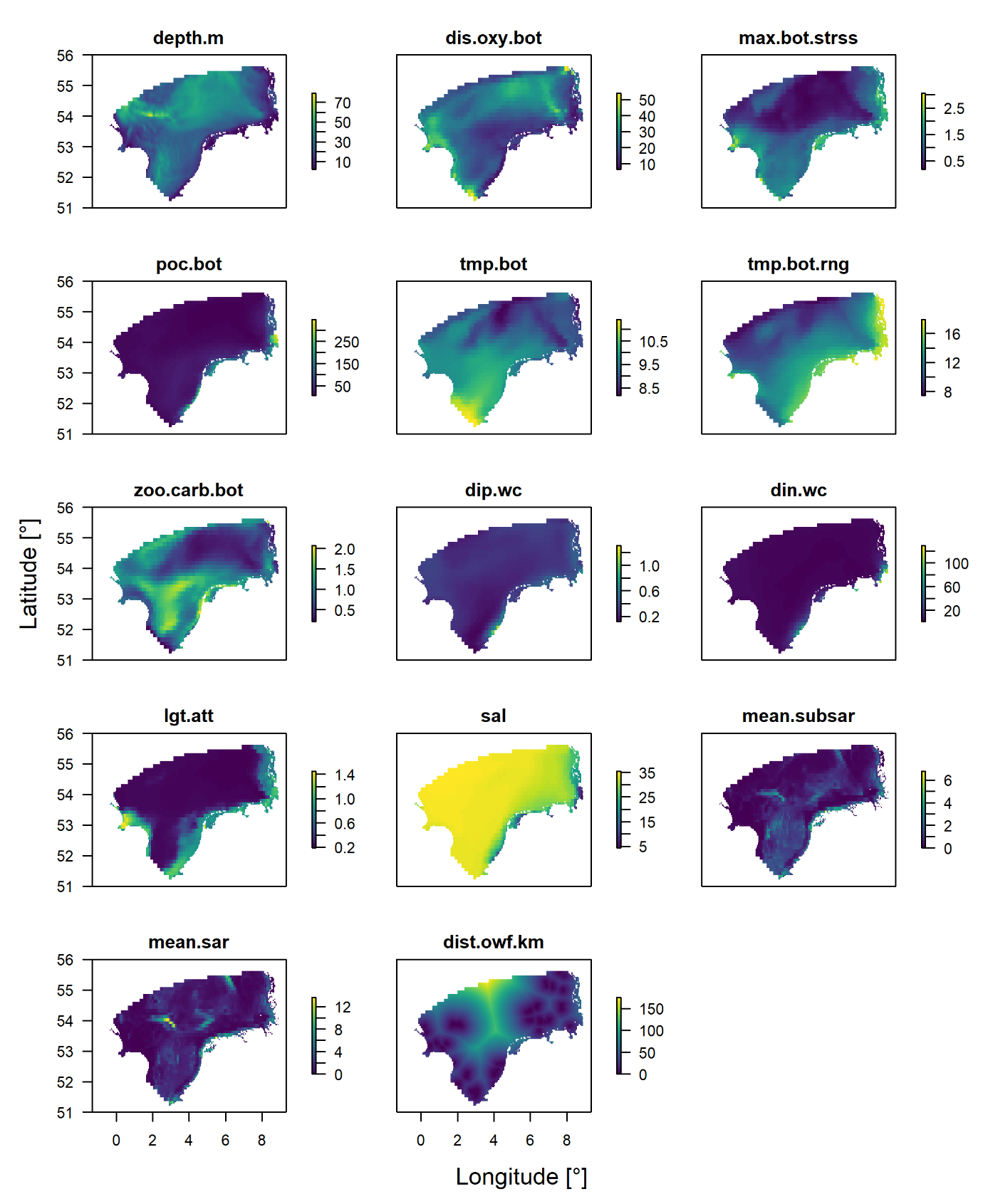

| **Table S2. Overview on predictors to model species distribution.** | | | | | |
| --- | --- | --- | --- | --- | --- |
| **Predictor** | **Description** | **Temporal resolution** | **Depth layer** | **Unit** | **Source** |
| Depth.m | Depth | Constant | Bottom layer | m | Hereon MOSSCO-GETM model |
| dis.oxy.bot | Dissolved oxygen | Mean of monthly means from 2004 - 2012 | Bottom layer | mmol O_2_/m^3^ | Hereon MOSSCO-GETM model |
| max.bot.strss | Shear stress | Mean of monthly means from 2004 - 2012 | Bottom layer | Pa | Hereon MOSSCO-GETM model |
| poc.bot | Particulate organic matter in the soil? | Mean of monthly means from 2004 - 2012 | average | mmol C/m3 | Hereon MOSSCO-GETM model |
| tmp.bot | Water temperature | Mean of monthly means from 2004 - 2012 | Bottom layer | °C | Hereon MOSSCO-GETM model |
| tmp.rng.bot | Range of watertemperature | Range of monthly means from 2004 - 2012 | Bottom layer | °C | Hereon MOSSCO-GETM model |
| zoo.carb.bot | Zooplankton | Mean of monthly means from 2004 - 2012 | Bottom layer | mmol C/m3 | Hereon MOSSCO-GETM model |
| dip.wc | Dissolved inorganic phosphorus | Mean of monthly means from 2004 - 2012 | Depth-average | mmol P/m3 | Hereon MOSSCO-GETM model |
| din.wc | Dissolved inorganic nitrogen | Mean of monthly means from 2004 - 2012 | Depth-average | mmol N/m3 | Hereon MOSSCO-GETM model |
| lgt.att | Light attenuation | Mean of monthly means from 2004 - 2012 | Water column |  | Hereon MOSSCO-GETM model |
| sal | Salinity | Mean of monthly means from 2004 - 2012 | Water column |  |  |
| mean.sar | Swept area ratio surface (reflects fishing pressure of mostly otter board trawls) | Mean of annual values from 2009 to 2020 | Sea floor | Proportion of grid cell area (values >1 indicate grid cells were swept area is larger than grid cell area) | ICES 2021 |
| mean.subsar | Swept area ratio subsurface (reflects fishing pressure of mostly beam trawls) | Mean of annual values from 2009 to 2020 | See flor | Proportion of grid cell area | ICES 2021 |
| dist.owf.km | Distance to constructed offshore wind farms | In operation or under construction until 2022 | Water volumn | km | 4Coffshore |

**Figure S3.** Temperature variables i.e. sea bottom temperature (tmp.bot) and sea bottom temperature range (tmp.bot.rng) for the three future scenarios (2030-2040, 2060-2070, 2090-2100).

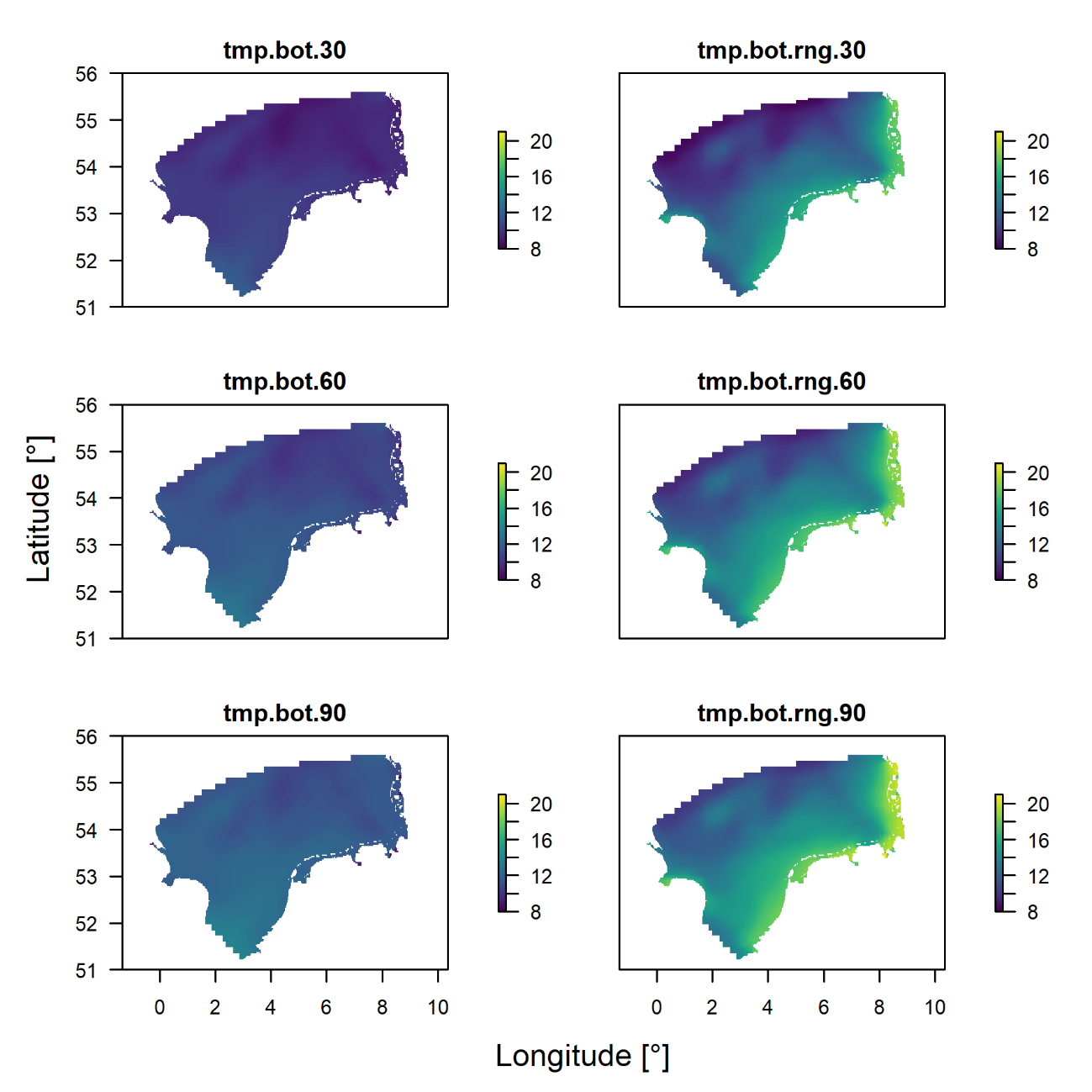

**Figure S4.** Scree-plot to determine number of clusters in K-means clustering of the demersal fish community. The red dot indicates the number of K after which the sum-of-squares error within clusters (WSS) declines to less than 66 % of the initial WSS at K = 1 (blue dashed line), defining the ‘knee’ of the plot (at K = 6).

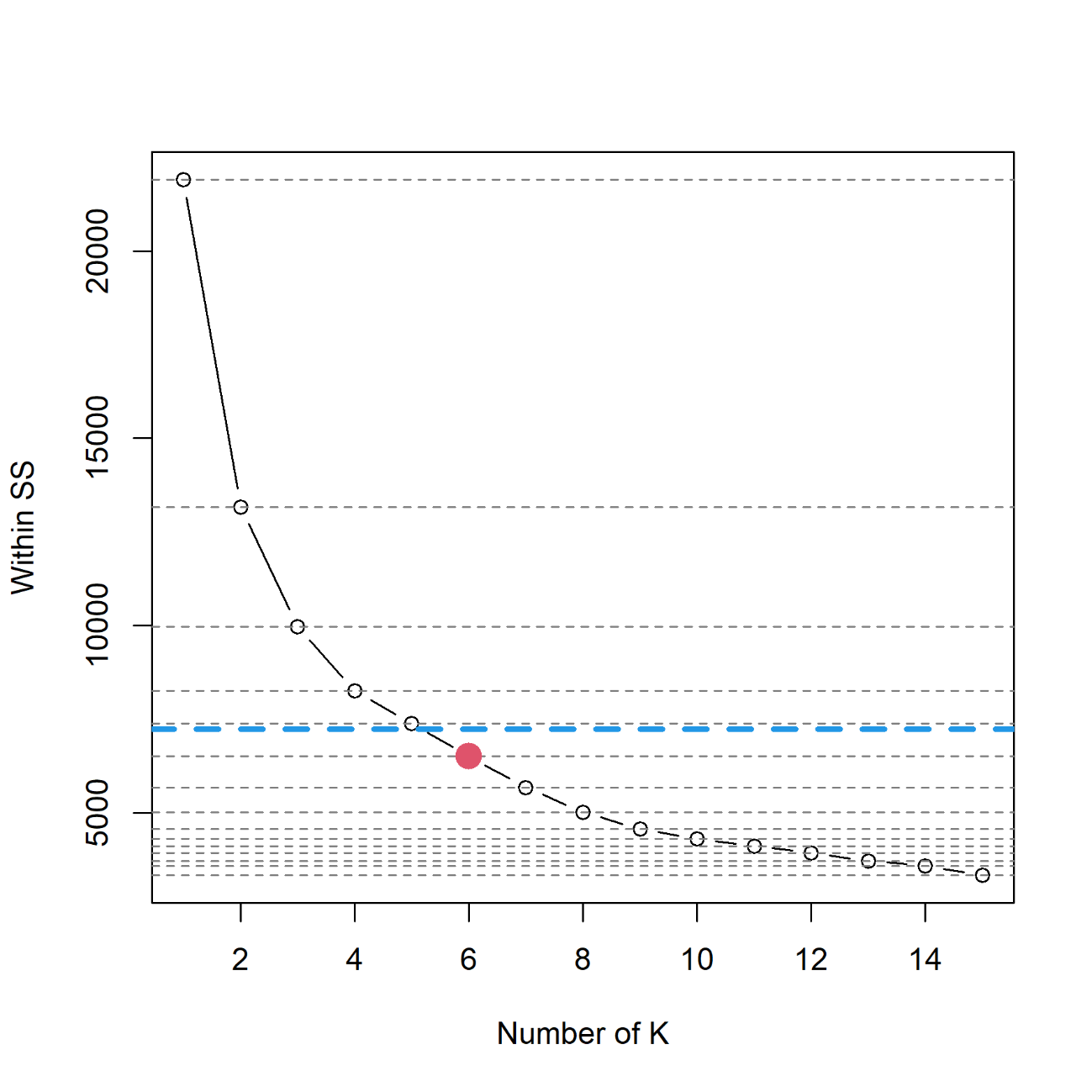

**Figure S5.** Performance metrics of SDM models for probability-of-occurrence (POC) and biomass. Blue colours represent POC related metrics, red colours represent biomass related metrics

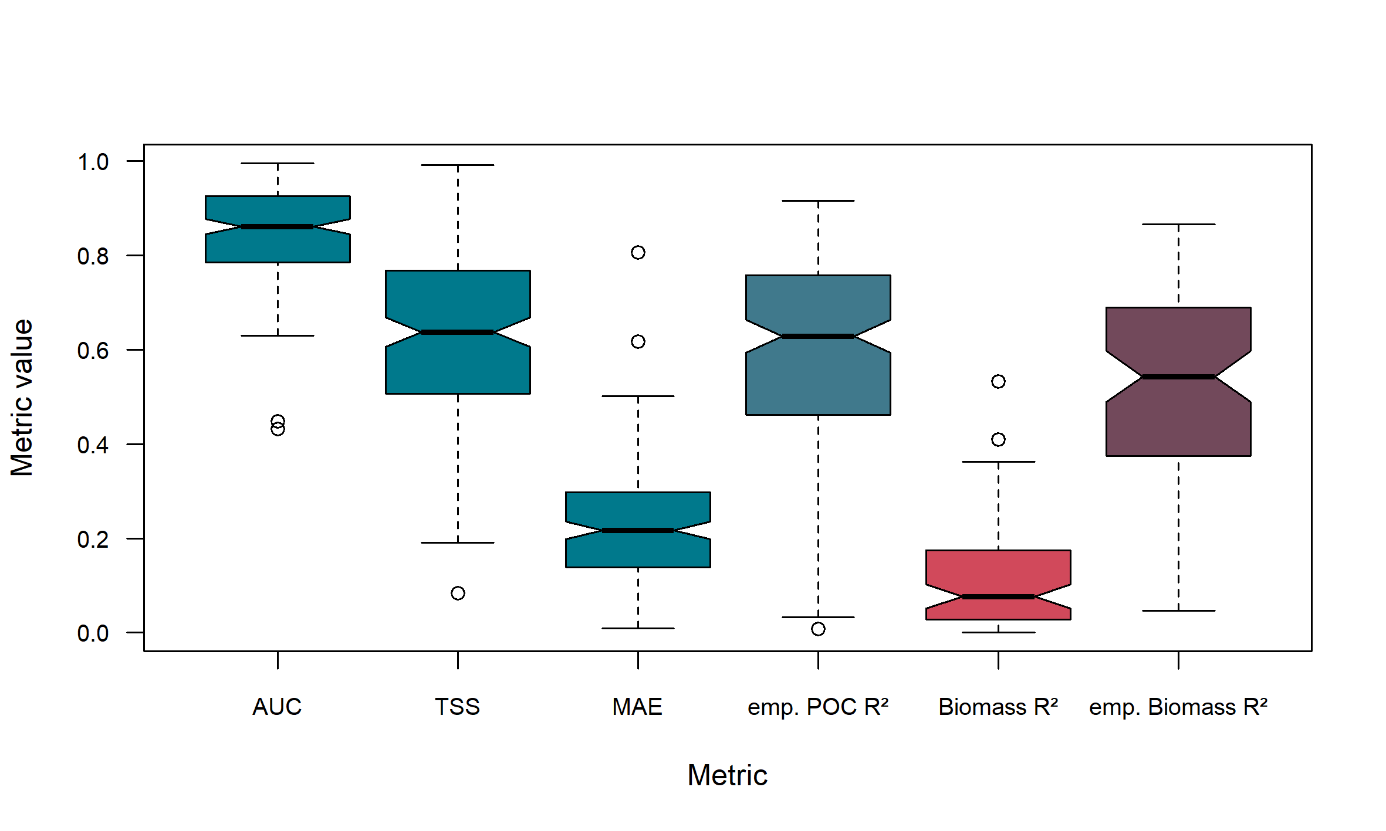

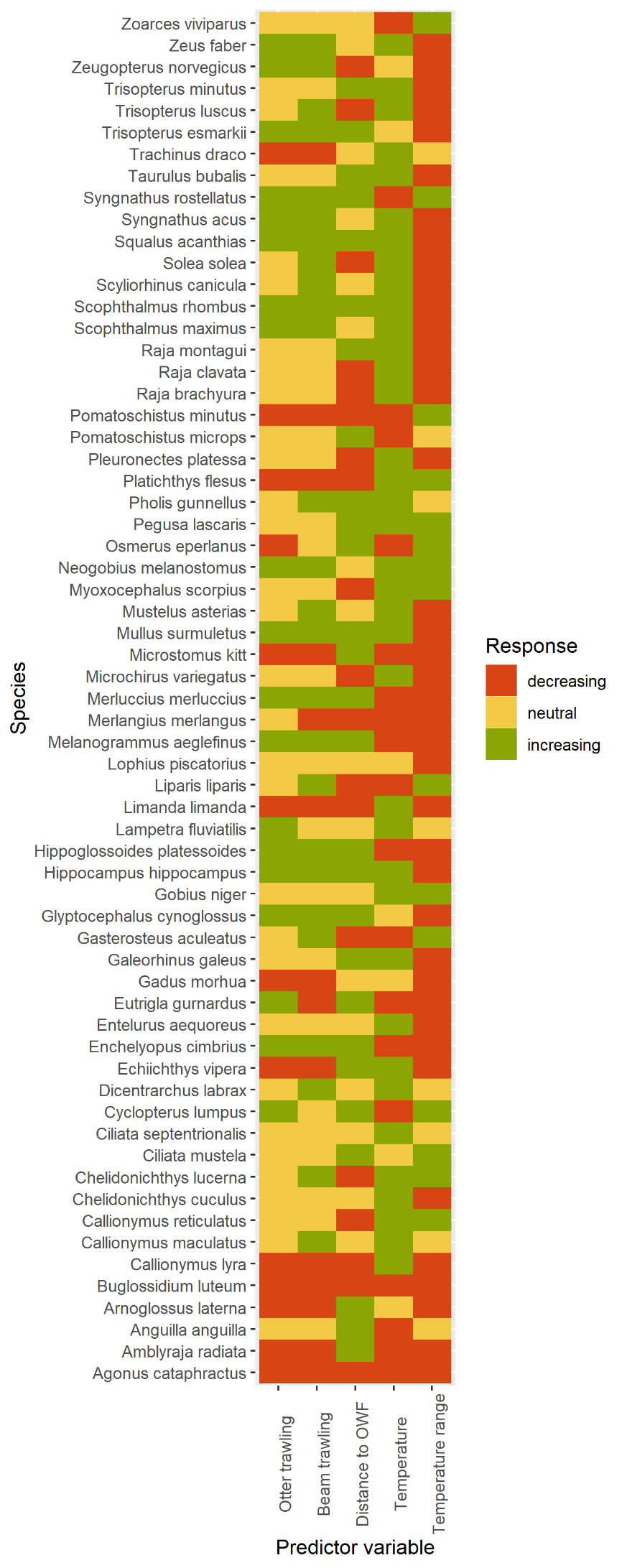

**Figure S6.** Species responses to human (otter trawling, beam trawling, distance to offshore wind parks) and environmental pressures (bottom temperature and bottom temperature range). Colours refer to significant negative (red) or positive slopes (green) of generalized linear models fitted to partial dependence data. In case the partial dependence data did not result in a significant GLM, the dependency was classified as neutral (yellow).

**Figure S7.** Changes in the distribution of red gurnard *Chelidonichtys cuculus* from present (2014-2023) until 2100 shown as probability-of-occurrence (POC, left) and biomass (kg km^-2^, right).

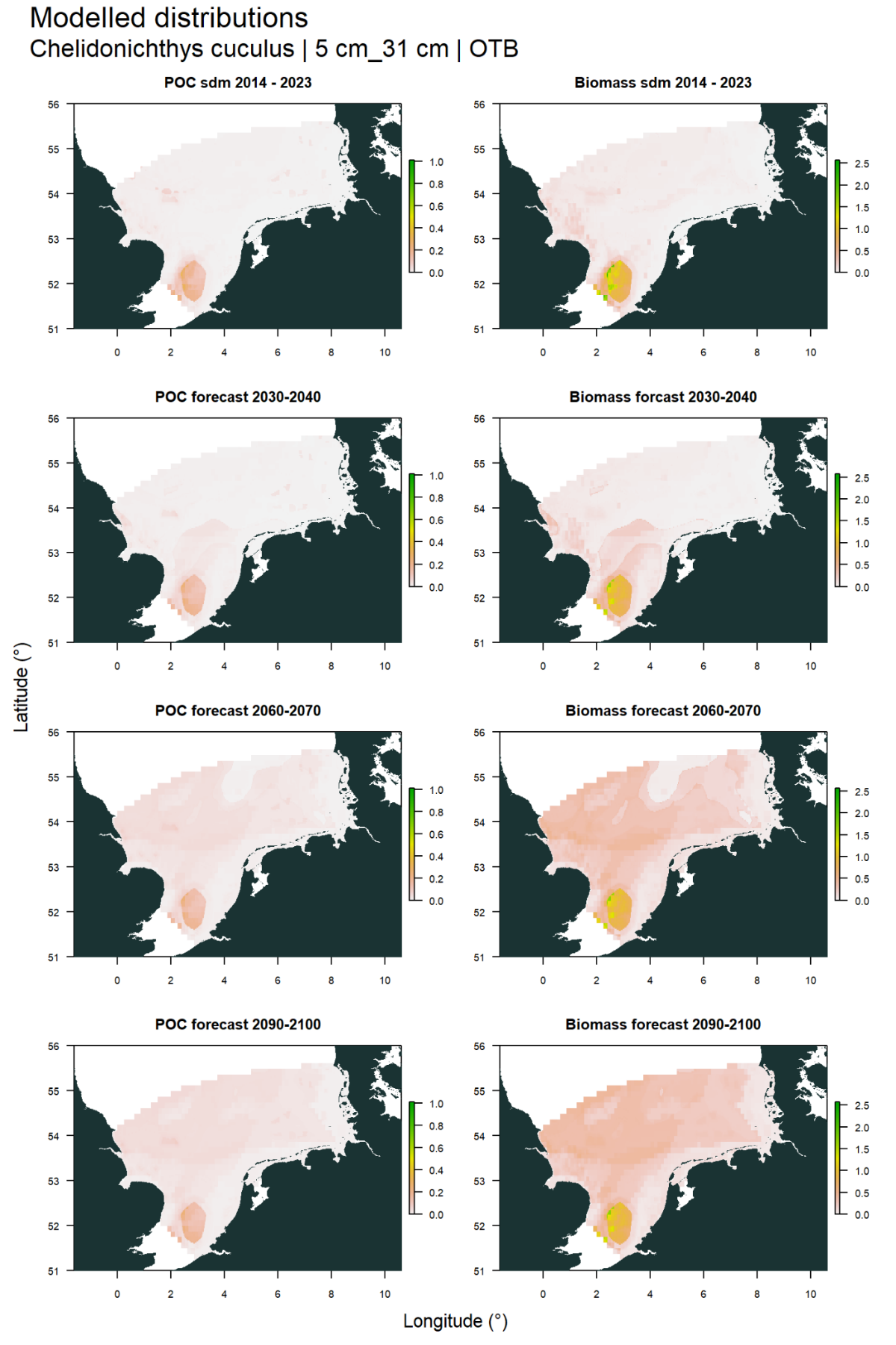

**Figure S8.** Co-occurrence of core areas (dark blue, based on biomass) of commercially relevant taxa and offshore windfarms (red). Note that even though 70 species were considered as commercially relevant species (Table S1), for only 52 species biomass data was available.

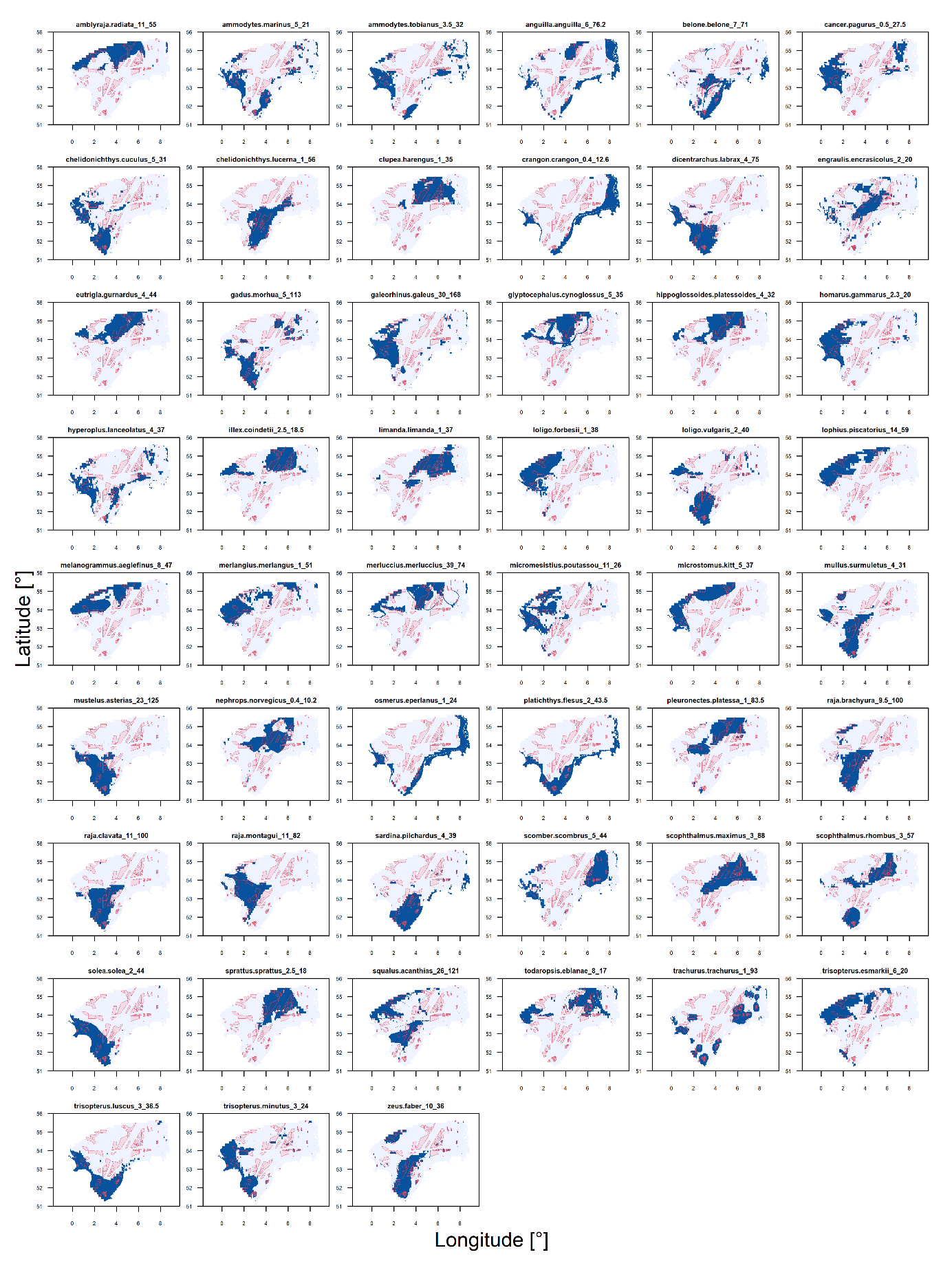

**Figure S9.** Overlap between core areas (based on probability of occurrence) of fish of conservation concern and offshore wind farms (blue) and marine protected areas (red).

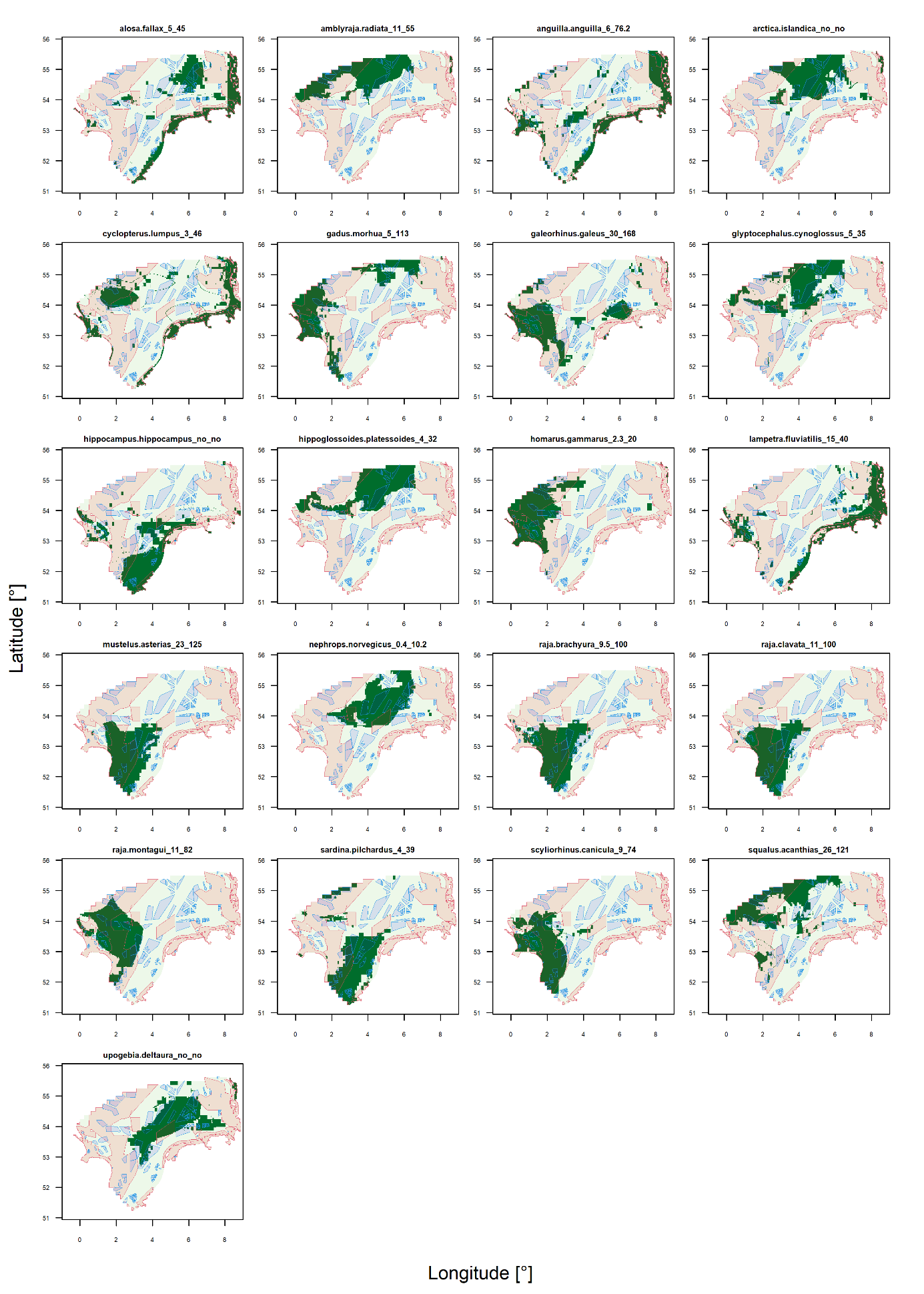
